# Chromosome-length genome assembly and structural variations of the primal Basenji dog (*Canis lupus familiaris*) genome

**DOI:** 10.1101/2020.11.11.379073

**Authors:** Richard J. Edwards, Matt A. Field, James M. Ferguson, Olga Dudchenko, Jens Keilwagen, Benjamin D. Rosen, Gary S. Johnson, Edward S. Rice, LaDeanna Hillier, Jillian M. Hammond, Samuel G. Towarnicki, Arina Omer, Ruqayya Khan, Ksenia Skvortsova, Ozren Bogdanovic, Robert A. Zammit, Erez Lieberman Aiden, Wesley C. Warren, J. William O. Ballard

## Abstract

**Background:** Basenjis are considered an ancient dog breed of central African origins that still live and hunt with tribesmen in the African Congo. Nicknamed the barkless dog, Basenjis possess unique phylogeny, geographical origins and traits, making their genome structure of great interest. The increasing number of available canid reference genomes allows us to examine the impact the choice of reference genome makes with regard to reference genome quality and breed relatedness.

**Results:** Here, we report two high quality *de novo* Basenji genome assemblies: a female, China (CanFam_Bas), and a male, Wags. We conduct pairwise comparisons and report structural variations between assembled genomes of three dog breeds: Basenji (CanFam_Bas), Boxer (CanFam3.1) and German Shepherd Dog (GSD) (CanFam_GSD). CanFam_Bas is superior to CanFam3.1 in terms of genome contiguity and comparable overall to the high quality CanFam_GSD assembly. By aligning short read data from 58 representative dog breeds to three reference genomes, we demonstrate how the choice of reference genome significantly impacts both read mapping and variant detection.

**Conclusions:** The growing number of high-quality canid reference genomes means the choice of reference genome is an increasingly critical decision in subsequent canid variant analyses. The basal position of the Basenji makes it suitable for variant analysis for targeted applications of specific dog breeds. However, we believe more comprehensive analyses across the entire family of canids is more suited to a pangenome approach. Collectively this work highlights the importance the choice of reference genome makes in all variation studies.

## Background

Dogs were the first animals to be domesticated by humans some 30,000 years ago [1] and exhibit exceptional levels of breed variation as a result of extensive artificial trait selection [2]. It is not clear whether dogs were domesticated once or several times, though the weight of accumulating evidence suggests multiple events [3–9]. By establishing genome resources for more ancient breeds of dog, we can explore genetic adaptations perhaps unique to the modern dog breeds. Basenjis are an ancient breed that sits at the base of the currently accepted dog phylogeny [10]. Basenji-like dogs are depicted in drawings and models dating back to the Twelfth Dynasty of Egypt [11] and they share many unique traits with pariah dog types. Like dingoes and New Guinea Singing dogs (NGSD), Basenjis come into oestrus annually—as compared to most other dog breeds, which have two or more breeding seasons every year. Basenjis, dingoes and NGSDs are prone to howls, yodels, and other vocalizations over the characteristic bark of modern dog breeds. One explanation for the unusual vocalisation of the Basenji is that the larynx is flattened [12]. The shape of the dingo and NGSD larynx is not reported.

Basenjis were originally indigenous to central Africa, wherever there was tropical forest. Primarily, what is now the DRC Congo, Southern Sudan, Central African Republic and the small countries on the central Atlantic coast. Today their territory has shrunk to the more remote parts of central Africa. The Basenji probably made its debut in the western world in around 1843. In a painting of three dogs belonging to Queen Victoria and Prince Albert entitled “Esquimaux, Niger and Neptune”, Niger is clearly a Basenji. In total, 71 Basenjis have been exported from Africa and, to date, ∼ 56 have been incorporated into the registered Basenji breeding population.

The first dog genome to be sequenced was of Tasha the Boxer [13], which was a tremendous advance and continues to be the resource guiding canine genomics research today. The Boxer is a highly derived brachycephalic breed that has been subjected to generations of artificial selection. Further, due to its discontiguous sequence representation it has been difficult to accurately detect structural variations (SVs) in other domestic dog breeds. Now, a new generation of breed-specific chromosome-level genome reference assemblies are becoming available (5 breeds in October 2020 according to the NCBI assembly archive). For example, we previously published a chromosome-level German Shepherd dog (GSD) genome assembly (CanFam_GSD) that is comprised of only 410 scaffolds and 716 contigs [14].

Here, we first report the sequence and *de novo* assembly of two Basenji genomes, female and male. We then compare these assemblies with the Boxer (CanFam3.1) [15] and GSD (CanFam_GSD) [14]. We conduct pairwise comparisons and report single-nucleotide variants (SNVs) and SVs between Basenji, Boxer and GSD. We distinguish an SNV as a variation in a single nucleotide without any limitations on its frequency. SV comprises a major portion of genetic diversity and its biological impact is highly variable. Chen et al. [16] used high-resolution array comparative genome hybridization to create the first map of DNA copy number variation (CNV) in dogs. Many canine CNVs were shown to effect protein coding genes, including disease and candidate disease genes, and are thus likely to be functional. In this study, we find all types of genetic variation are impacted by the choice of reference genome. The basal position of the Basenji makes it useful as a general reference for variant analysis, but the generation of clade-specific genomes is likely to be important for canine nutrition and disease studies. We recommend a pan-genome approach for comprehensive analyses of canid variation.

## Results

### Basenji female assembly, CanFam_Bas

The female Basenji, China (Fig. 1A), was initially assembled from 84.5 Gb Oxford Nanopore Technologies (ONT) PromethION reads (approx. 35x depth based on a 2.41 Gb genome size) using Flye (v2.6) [17, 18] and subjected to long read polishing with Racon v1.3.3 [19] and Medaka 1.03 [20] (Supplementary Fig 1A, Additional File 1). Additional short read Pilon [21] error-correction was performed with 115.1 Gb (approx. 47.7x) BGIseq data. Hi-C proximity ligation was used with the DNA zoo pipeline [22–24] to scaffold 1,657 contigs into 1,456 scaffolds, increasing the N50 from 26.3 Mb to 63.1 Mb and decreasing the L50 from 33 to 14 (Figs 2 and 3, Supplementary Table 1, Additional File 2). Scaffolds were gap-filled by PBJelly (pbsuite v.15.8.24) [25] using the ONT data, reducing the number of gaps from 348 to 148 and the number of scaffolds to 1,407. Following a final round of Pilon [21] BGIseq-polishing, scaffolds were mapped onto the CanFam3.1 [13] using PAFScaff v0.40 [14, 26]. Diploidocus v0.9.0 vector filtering [27] removed one 5.7kb contig and masked a 3.3kb region of Chromosome X as lambda phage (J02459.1) contamination. Seven rounds of iterative Diploidocus tidying of the remaining sequences removed 277 (832 kb) as low coverage/quality and 481 (1.58 Mb) as probable haplotigs, retaining 483 core scaffolds and 165 probable repeat-heavy sequences [14] as China v1.0 (Fig 3, Supplementary Fig 2, Additional File 1, Supplementary Table 2, Additional File 2).

**Figure 1.**
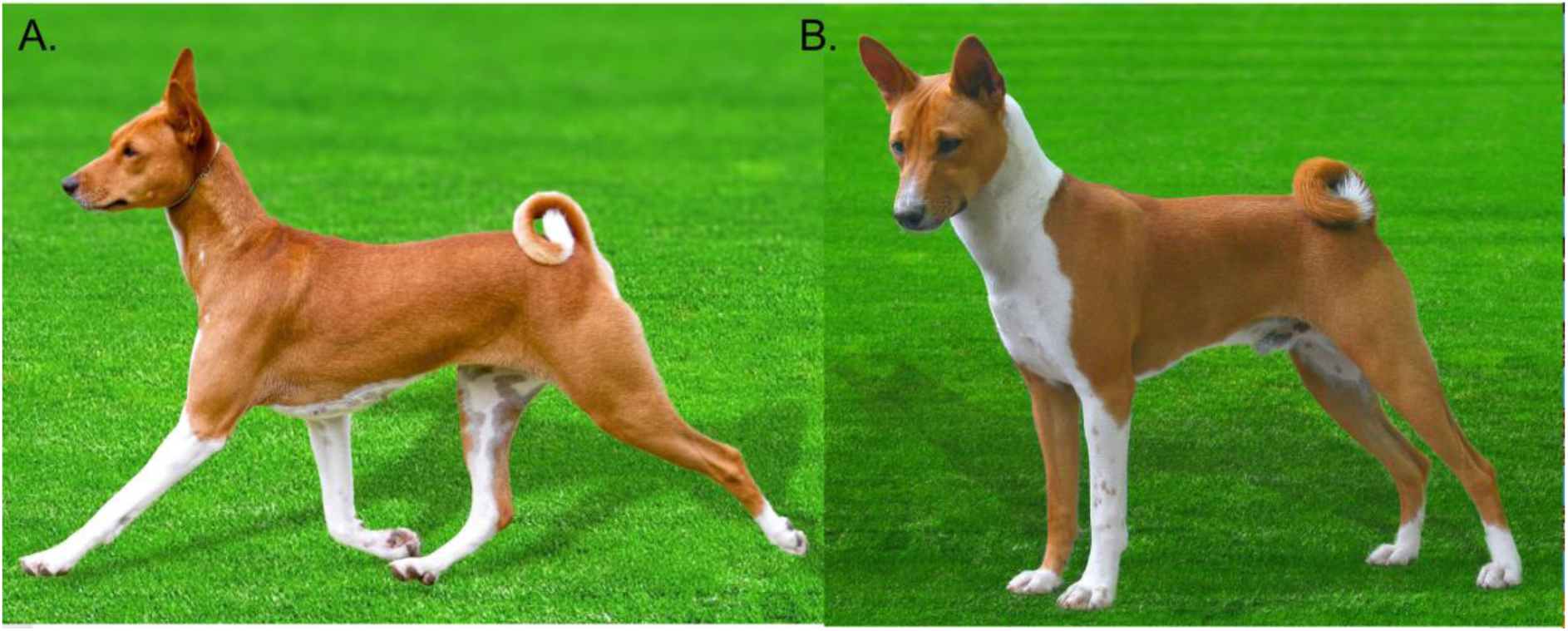
The Basenji dogs included in the study. **A**. China. Is registered as Australian Kennel Club Supreme Champion Zanzipow Bowies China Girl. Her registration is #2100018945. She was born in 2016 and she is free of all known genetic diseases. Her sire and dam are Australian bred and her most recent ancestor from Africa was 18 generations ago. *Photo credit: Dylan Edgar.* **B**. Wags. Is registered as American Kennel Club Champion Kibushi the Oracle, born in in 2008. His registration number is HP345321/01. His sire is an American bred dog while his dam was imported from the Haut-Ule district of the DRC Congo, 3°24’04.0”N 27°19’04.6”E, in 2006. *Photo credit: Jon Curby*.

**Figure 2.**
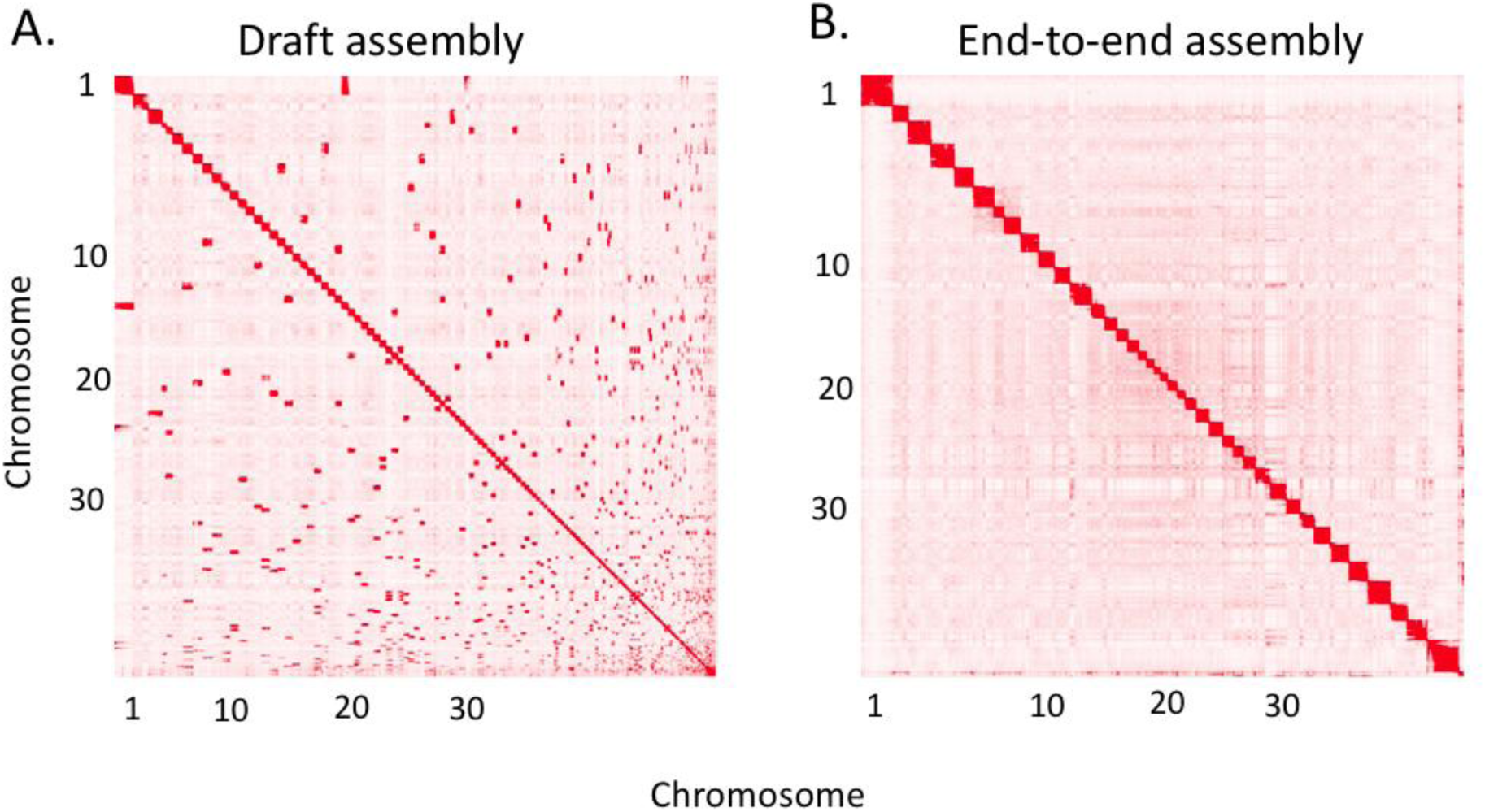
Contact matrices generated by aligning the CanFam_Bas (China) Hi-C data set to the genome assembly. **A**. before the Hi-C upgrade (draft assembly). **B.** After Hi-C scaffolding (End-to-end assembly).

**Figure 3.**
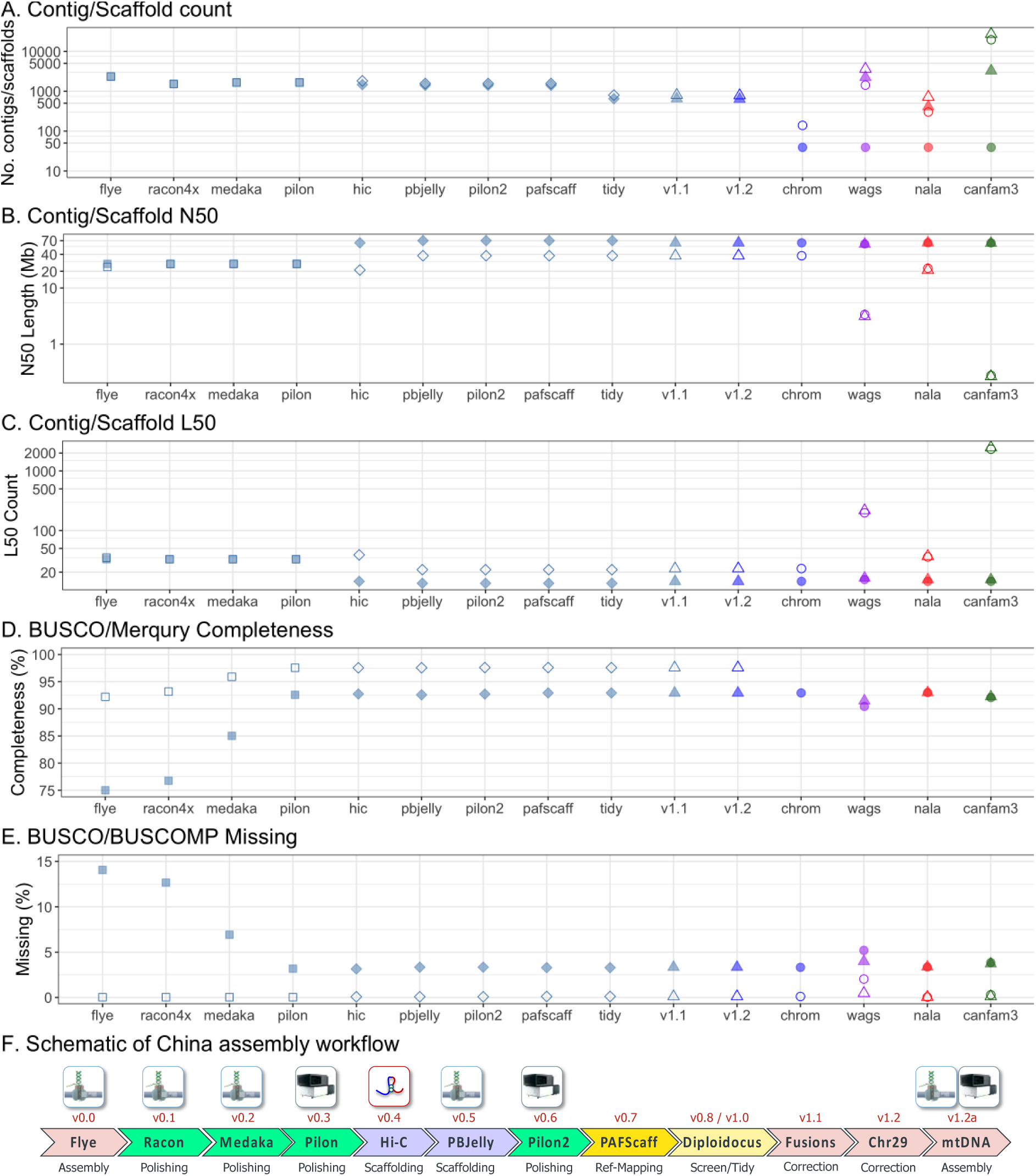
Key contiguity, quality and completeness metrics for different assembly stages and comparison dog genomes. Square, pre-scaffolded China; Diamond, scaffolded China; Triangle, complete assembly; Circle, main chromosome scaffolds only; Blue, China; Purple, Wags; Red, CanFam_GSD; Green, CanFam3.1. **A.** Contig (open) and scaffold (filled) numbers. **B.** Contig (open) and scaffold (filled) N50. **C.** Contig (open) and scaffold (filled) L50. **D.** Genome completeness estimated by BUSCO v3 (filled) and Merqury (open). **E.** The percentage of missing BUSCO genes (filled) and BUSCOMP genes (those found to be Complete in any assembly) (open). **F.** Schematic of China assembly workflow. CanFam_Bas is China v1.2.

#### Genome assembly correction

Two pairs of fused chromosomes in China v1.0 were incorrectly joined by PBJelly. Pre-gap-filled HiC scaffolds were mapped onto the assembly using Minimap2 v2.17 [28] and parsed with GABLAM v2.30.5 [29] to identify the gap corresponding to the fusion regions. These were manually separated into four individual chromosomes, gap lengths standardised, and scaffolds re-mapped onto CanFam3.1 using PAFScaff v0.4.0. D-GENIES [30] analysis against CanFam_GSD chromosomes confirmed that PBJelly had incorrectly fused two pairs of chromosomes: chromosomes 8 with 13, and chromosome 18 with 30. These were manually separated and the assembly re-mapped onto CanFam3.1 as China v1.1. PAFScaff assigned 112 scaffolds to chromosomes, including 39 nuclear chromosome-length scaffolds.

It was observed that the mitochondrial chromosome was missing and China v1.1 Chromosome 29 contained a 33.2 kb region consisting of almost two complete copies of the mitochondrial genome that were not found in other dog genome assemblies. The 26 ONT reads that mapped onto both flanking regions were reassembled with Flye v.2.7.1 [17, 18] into a 77.2 kb chromosome fragment, which was polished with Racon v1.3.3 [19] and Medaka 1.03 [20]. This was mapped back on to the Chromosome 29 scaffold with GABLAM v2.30.5 [29] (blast+ v2.9.0 megablast [31, 32]) and the mitochondrial artefact replaced with the corrected nuclear mitochondrial DNA (NUMT) sequence. Finally, scaffolds under 1 kb were removed to produce the China v1.2 nuclear genome that we name CanFam_Bas.

#### Mitochondrial genome assembly

In total, 4,740 ONT reads (52.1 Mb) mapping on to mtDNA were extracted. To avoid NUMT contaminants, a subset of 80 reads (1.32 Mb) with 99%+ mtDNA assignment and 99%+ mtDNA coverage, ranging in size from 16,197 kb to 17,567 kb, were assembled with Flye 2.7b-b1526 [17, 18] (genome size 16.7 kb) into a 33,349 bp circular contig consisting of two mtDNA copies. This contig was polished with Racon [19] and Medaka [20], before being re-circularised to a single-copy matching the CanFam3.1 mtDNA start position. After final Pilon [21] correction of SNPs and indels, the 16,761 bp mitochondrial genome was added to the CanFam_Bas assembly.

#### CanFam_Bas (China) reference genome

The resulting chromosome-length CanFam_Bas reference genome is 2,345,002,994 bp on 632 scaffolds with 149 gaps (76,431 bp gaps) (Table 1). The 39 nuclear plus mitochondrial chromosome scaffolds account for 99.3% of the assembly and show a high level of synteny with CanFam3.1 and CanFam_GSD (Fig 4). CanFam_Bas represents the most contiguous dog chromosomes to date, with a contig N50 of 37.8 Mb and contig L50 of 23, which is slight improvement over CanFam_GSD and considerably more contiguous than the standard dog reference genome, CanFam3.1 (Fig 3, Table 1). The completeness and accuracy of the genome as measured by BUSCO v3 [33] (laurasiatherian, *n*=6253) is also superior to CanFam3.1 and approaches that of CanFam_GSD (92.9% Complete, 3.75% Fragmented, 3.34% Missing).

**Figure 4.**
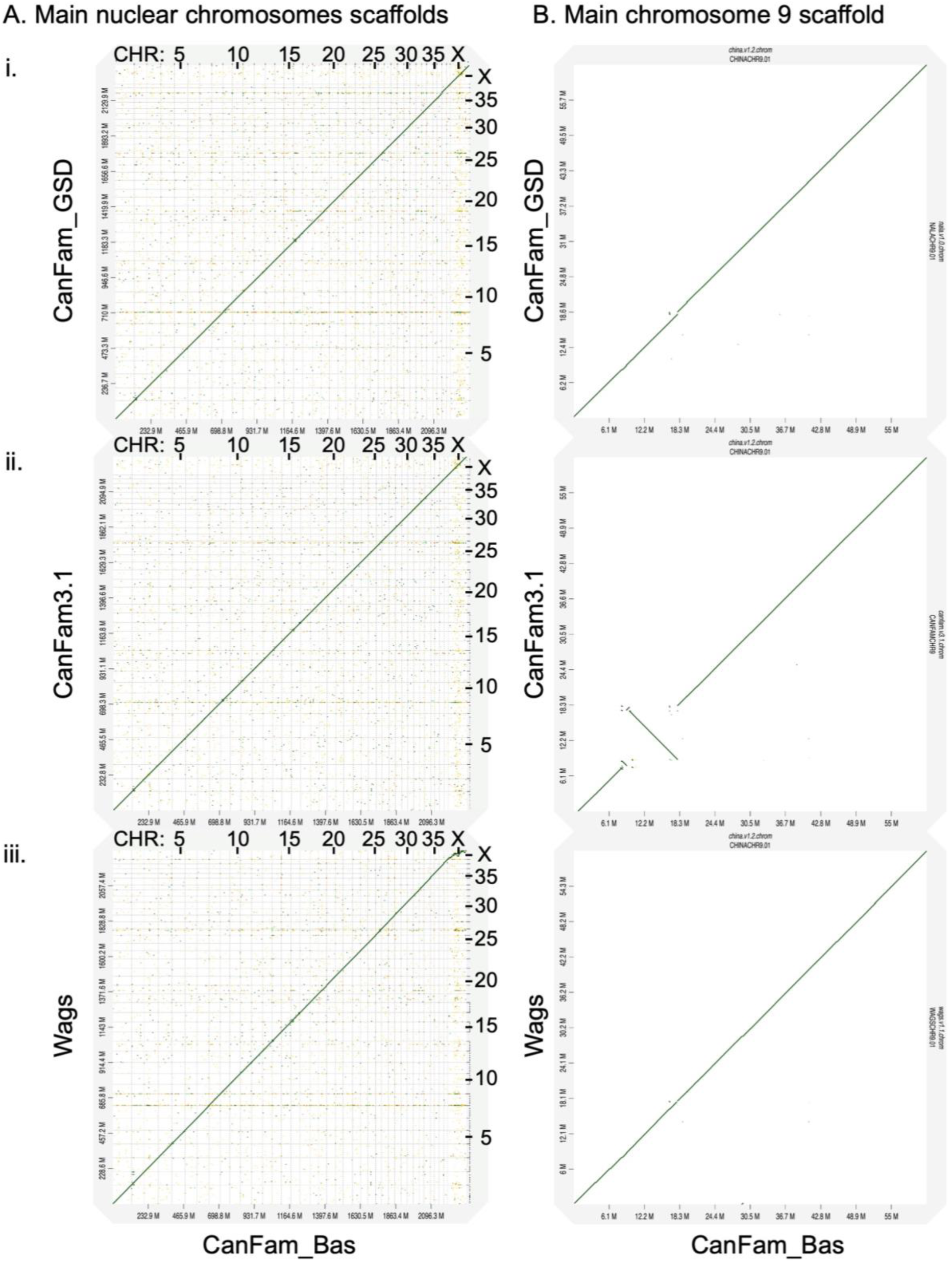
D-GENIES synteny plots of main chromosome scaffolds for three dog genome assemblies against CanFam_Bas. In each case, CanFam_Bas (China v1.2) is on the x-axis and the comparison assembly on the y-axis. Gridlines demarcate scaffolds. Thick black lines indicate regions of genomic alignment. **A**. All-by-all main chromosome scaffold alignments with **(i)** CanFam_GSD, **(ii)** CanFam_3.1, and **(iii)** Wags. **B**. Main chromosome 9 scaffold alignment with **(i)** CanFam_GSD, **(ii)** CanFam_3.1, and **(iii)** Wags.

**Table 1.**
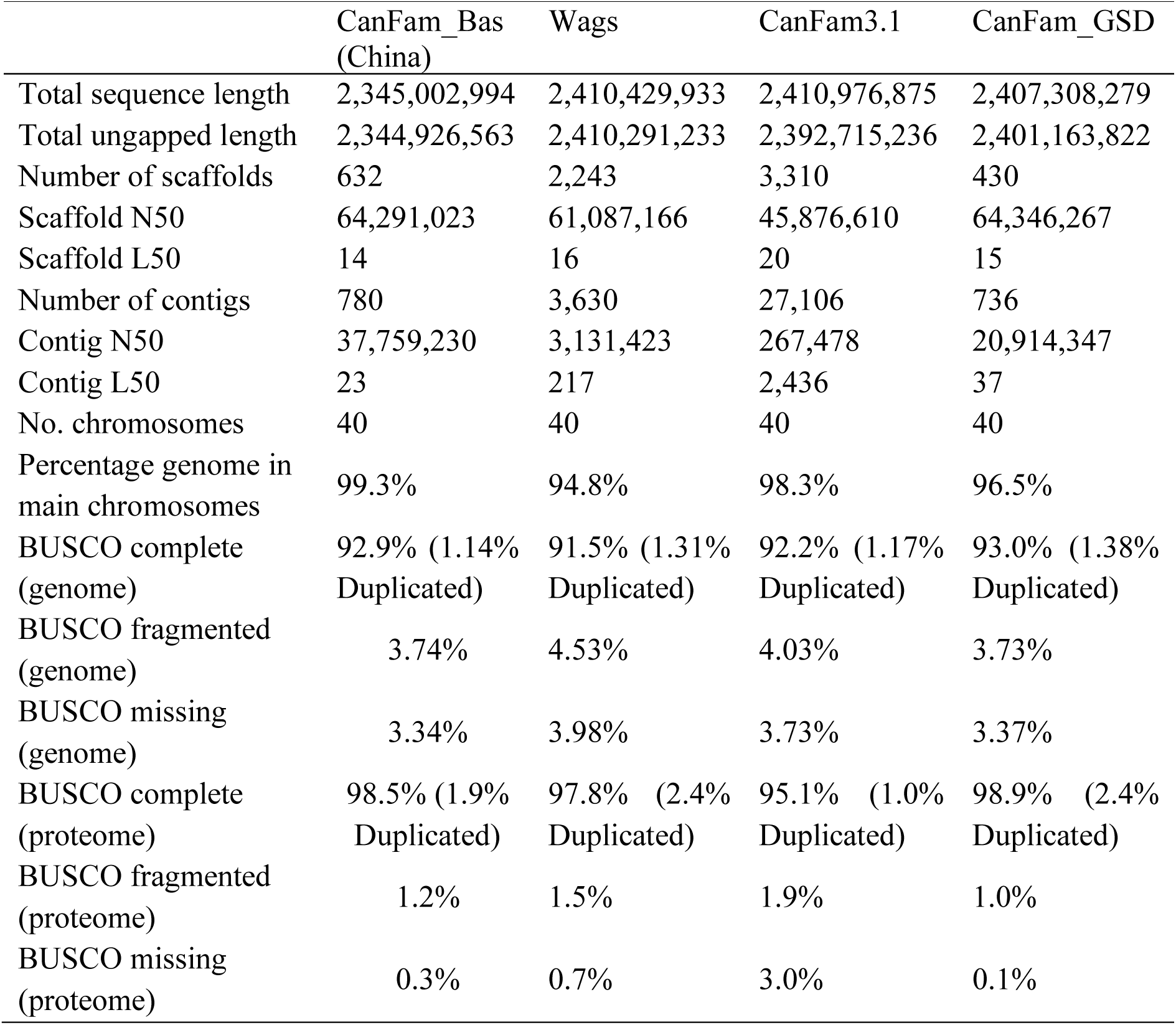
Genome assembly and annotation statistics for Basenji assemblies vs CanFam3.1 and CanFam_GSD.

### Methylomic identification of putative regulatory elements

Additionally, we profiled whole genome methylation of Basenji’s blood DNA using MethylC-seq [34]. Numbers of unmethylated and highly methylated CpG sites in Basenji’s genome were similar to that of GSD (Supplementary Fig. 3A, Additional File 1). Importantly, high resolution DNA methylation data can be utilised to identify the putative regulatory elements in a given tissue type. That is, CpG-rich unmethylated regions (UMRs) mostly correspond to gene promoters, while CpG-poor low-methylated regions (LMRs) correspond to distal regulatory elements, both characterised by DNAse I hypersensitivity [35]. Using MethylSeekR algorithm [36], we performed the segmentation of Basenji DNA methylome and discovered 20,660 UMRs and 54,807 LMRs (Supplementary Fig. 3B,C, Additional File 1), in line with previously reported numbers in mammalian genomes [14, 36, 37]. Genome-wide and locus-specific CpG methylation called by MethylC-seq correlated strongly with that called directly from the ONT data (Supplementary Fig. 3D-F, Additional File 1), confirming the robustness of determined DNA methylation profile of the blood DNA.

### Male Basenji assembly, Wags

For the male Basenji, Wags, (Fig 1B) we generated Pacific Bioscience Single Molecule Real Time (SMRT) sequences to approximately 45x genome coverage and assembled the genome to ungapped size of 2.41 Gb (Supplementary Fig 1B, Additional File 1). Assembly contiguity metrics of 3,630 total contigs show N50 contig and scaffold lengths of 3.1 and 61 Mb length, respectively (Table 1). Wags alignment to China revealed a high level of synteny. However, the Wags assembly of the X chromosome is smaller in size (59 Mb vs 125 Mb) and shows multiple rearrangements as a result of lower sequence coverage on the sex chromosomes (∼21x). We were unable to accurately place 124.4 Mb of Wags sequence on 2,204 scaffolds (2,210 contigs), including 651 contigs with a total length of 45.6 Mb mapped on to the CanFam3.1 X chromosome by PAFScaff. Therefore, all comparative analyses reported herein were done with CanFam_Bas. In addition, the Wags assembly includes 3.6 Mb of the Basenji dog Y for future comparative studies of this unique chromosome.

### Genome annotation

The CanFam_Bas and Wags assemblies were annotated using the homology-based gene prediction program GeMoMa v1.6.2beta [38] and nine reference species [14]. In total, CanFam_Bas and Wags had similar numbers of predicted protein-coding genes at 27,129 (68,251 transcripts) and 27,783 (65,511) transcripts, respectively (Supplementary Table 3, Additional File 2). Analysing the longest protein isoform per gene with BUSCO v3 [33] (laurasiatherian, *n*=6253, proteins mode), CanFam_Bas was measured to be 98.5 % complete (1.9% duplicated) and Wags was measured as 97.8% complete (2.4% duplicated). Both proteomes compare favourably with CanFam3.1 in terms of completeness (Table 1). To correct for differences introduced by the annotation method, CanFam3.1 was annotated with the same GeMoMa pipeline. Approximately 90% of the Quest For Orthologues (QFO) reference dog proteome [39] is covered by each GeMoMa proteome, confirming comparable levels of completeness (Supplementary Table 3, Additional File 2).

When the CanFam_Bas GeMoMa proteome was compared to Wags, CanFam3.1 and CanFam_GSD, over 91% genes had reciprocal best hits for at least one protein isoform (Supplementary Table 3, Additional File 2). To investigate this further, the Wags, CanFam3.1 and CanFam_GSD genomes were mapped onto CanFam_Bas and the coverage for each gene calculated with Diploidocus v0.10.0. Of the 27,129 predicted genes, 26,354 (97.1%) are found at least 50% covered in all four dogs, whilst only 30 (0.11%) are completely unique to CanFam_Bas. In total, Wags is missing 302 predicted genes, CanFam_GSD is completely missing 95 predicted genes, and CanFam3.1 is missing 211 predicted genes (Table 2). A considerably greater proportion of the missing genes in Wags (64.2% versus 11.4% in CanFam3.1 and 15.8% in CanFam_GSD) were on the X chromosome. To test for artefacts due to assembly errors we mapped the long read data for Wags and CanFam_GSD onto CanFam_Bas. Only 7 of the 302 missing Wags genes (2.3%) had no long read coverage, whilst 21/95 (22.1%) of genes missing in CanFam_GSD were confirmed by an absence of mapped long reads.

**Table 2.** Predicted copy numbers for CanFam_Bas GeMoMa genes based on A. assembly mapping, and B. long read mapping.

| **A. Dog** | **Missing** | **Partial**<br>**(<50%)** | **Single**<br>**(1n)** | **Duplicate**<br>**(2n)** | **3n+** |
| --- | --- | --- | --- | --- | --- |
| CanFam_Bas | 0<br>(0) | 0<br>(0) | 27129<br>(25788) | 0<br>(0) | 0<br>(0) |
| CanFam_Wags | 302<br>(108) | 120<br>(58) | 26103<br>(25035) | 486<br>(472) | 118<br>(115) |
| CanFam3.1 | 211<br>(187) | 167<br>(161) | 26404<br>(25125) | 251<br>(223) | 96<br>(92) |
| CanFam_GSD | 95<br>(80) | 48<br>(42) | 26586<br>(25304) | 299<br>(266) | 101<br>(96) |

| **B. Data** | **0n** | **0.5n** | **1n** | **1.5n** | **2n** | **2.5n+** |
| --- | --- | --- | --- | --- | --- | --- |
| CanFam_Bas<br>(ONT) | 2<br>(2) | 1257<br>(1201) | 24116<br>(22954) | 1508<br>(1403) | 80<br>(68) | 166<br>(160) |
| CanFam_Wags<br>(PacBio) | 7<br>(2) | 4717<br>(3476) | 21140<br>(21049) | 1028<br>(1024) | 79<br>(79) | 158<br>(158) |
| CanFam_GSD<br>(ONT) | 21<br>(18) | 1893<br>(1795) | 22412<br>(21350) | 2503<br>(2349) | 109<br>(98) | 191<br>(178) |

Figures in brackets exclude predicted genes on X chromosome.

### Amylase copy number

Two copies of the *Amy2B* gene were identified in a tandem repeat on Chromosome 6 of the CanFam_Bas assembly. The single-copy read depth for CanFam_Bas, calculated as the modal read depth across single copy complete genes identified by BUSCO v3 [33], was estimated to be 34x. This was verified by BUSCO complete genes, which gave mean predicted copy numbers of 1.008±0.005 (95% C.I.) (Supplementary Fig 4, Additional File 1). The two complete *Amy2B* coding sequence copies had a mean depth of 97.5x, equating to 2.87N, or a total copy number estimate of 5.78N (2 x 97.5 / 34). The full CanFam_GSD *Amy2B* repeat region was also found in two complete copies with a mean depth of 98.1x, estimating 5.77 copies (2 x 98.1 / 34). Similar results were obtained restricting analysis to reads at least 5 kb (6.01 gene copies; 5.98 region copies) or 10 kb (6.18 gene copies; 6.04 region copies) to minimise repeat-based fluctuations in read depth. In contrast, droplet digital PCR (ddPCR) estimated that the Basenji China had 4.5 copies per individual chromosome. This slight difference suggests that the raw sequence data slightly overestimated copies or the ddPCR primers did not capture all the genetic variation.

Wags assembly has a single copy of the *Amy2B* region, which includes 90% of the *Amy2B* coding sequence (data not shown). Single-copy depth analysis of Wags estimates 4.97 (90% at 253.8x) and 5.51 (100% at 253.5x) copies of the *AMY2B* coding sequence and tandem repeat unit, respectively. To estimate copy number in other Basenji dogs, short read data was downloaded from SRA for 11 Basenjis (Supplementary Table 4, Additional File 2) and mapped onto CanFam_BAS. AMY2B copy number was estimated as the ratio of mean read depth per AMY2B gene versus the mean depth for the whole genome. Estimated copy numbers ranged from 3.93 to 6.79, with a mean of 5.25 (Supplementary Table 4, Additional File 2). The same analysis was performed on China BGI data, yielding an estimate of 4.38 copies, consistent with the ddPCR results.

### Nuclear mitochondrial DNA fragments

During the assembly of the female Basenji genome (China v1.0), the mitochondrial genome was erroneously assembled into a NUMT fragment on chromosome 29. A blastn search (e < 10-4 [40]) identified 291 putative NUMT fragments, ranging in size from 34 bp to 6,580 bp, forming 212 NUMT blocks from 34 bp to 16,577 bp (Supplementary Table 5,6, Additional File 2). Fragments total 190.5 kb (approx. 11.4 mtDNA copies) and span the entire mtDNA with reasonably even coverage and no obvious bias (Supplementary Fig. 5A, Additional File 1), except for low coverage in a region of the D-loop as has been previously reported in primates [41]. All 291 NUMT fragments are well-supported with at least 3 reads that span the entire NUMT plus at least 5 kb each side (Supplementary Table 5, Additional File 2). Only 1 NUMT was not found to be full-length in CanFam_GSD (Supplementary Fig. 5B, Additional File 1). An additional 26 NUMTs are partially covered in CanFam3.1 and 9 are entirely absent. Whilst this could represent a breed difference, 19 of the 35 additional incomplete NUMTs in CanFam3.1 (65.5%) are also incomplete in Wags, whilst Wags has a further ten incomplete NUMTs that are present in CanFam3.1 (Supplementary Table 5, Additional File 2). This is consistent with these regions being generally harder to assemble and/or align. Further analyses of these regions may provide insight into domestic dog genealogies.

### Whole genome assembly comparisons

To discover unique large-scale structural differences in assembled genomes of the three breeds – Basenji, German Shepard and Boxer – we performed pairwise alignments of CanFam_Bas, CanFam3.1 and CanFam_GSD. Overall, genome synteny was maintained and there were limited large scale genomic rearrangements observed (Fig 4A). There was, however, a large inversion in CanFam3.1 that was not present in CanFam_Bas or CanFam_GSD (Fig 4B). To investigate this further, we aligned CanFam_Bas against Wags (Fig 4A iii). As expected, there was no inversion on Chromosome 9 (Fig 4B iii), suggesting the inversion or perhaps an assembly error, occurs in CanFam3.1.

### Long read structural variant detection

To generate a conservative set of high-quality structural variants, the consensus of ONT and SMRT long read sequences data from both Basenji (China female ONT and Wags male SMRT) and GSD (Nala female ONT and SMRT [14]) was mapped onto Basenji (CanFam_BAS), Boxer (CanFam3.1) and GSD (CanFam_GSD) reference genomes (Supplementary Table 7, Additional File 2). One difference in the data sets is that both ONT and SMRT reads from GSD were sourced from the same individual dog whereas for Basenji the ONT and SMRT reads are from different individual Basenji dogs. These high-quality SVs were additionally annotated for their overlap to both genes and exons. Of these high-quality ONT/SMRT consensus SVs, the Basenji long reads overlap 814 CanFam3.1 exons and 568 CanFam_GSD exons while the GSD long reads overlap 825 CanFam3.1 exons and 495 CanFam_Bas exons. In total, these SVs represent 92.19 Mb, 97.21 Mb, and 78.99 Mb of deleted sequence and 4.11 Mb, 4.09 Mb, and 7.69 Mb of inserted sequence for the CanFam_Bas, CanFam3.1, and CanFam_GSD assemblies respectively.

To reduce the impact of small indels arising from local mis-assembly errors, the high-quality consensus SVs were further reduced to those over 100 bp in length (Fig 5). Overall, we observe similar number of total SV calls from the Basenji long reads relative to CanFam_GSD and the GSD long reads relative to CanFam_BAS. Both breeds had a substantially larger number of consensus SV calls against the CanFam3.1 reference, with Basenji long reads generating more SVs calls than GSD long reads. We next overlapped the CanFam SV calls relative to CanFam3.1 and found 18,063 long read deletion calls overlapped between Basenji and GSD. For Basenji this represented 70.00% of the total 25,260 Basenji deletions while for GSD this represented 73.25% of the total 24,138 GSD deletions (Fig. 5B). Insertions were fewer in number and degree of overlap, however we still found 5,146 overlapping insertions between Basenji and GSD long reads representing 33.33% of the total 15,434 Basenji insertions and 36.46% of the total 14,111 GSD insertions (Fig. 5C). The high degree of overlap in SVs from GSD and Basenji relative to CanFam3.1 represent Boxer-specific SVs or potential issues with the current canid reference assembly.

**Figure 5.**
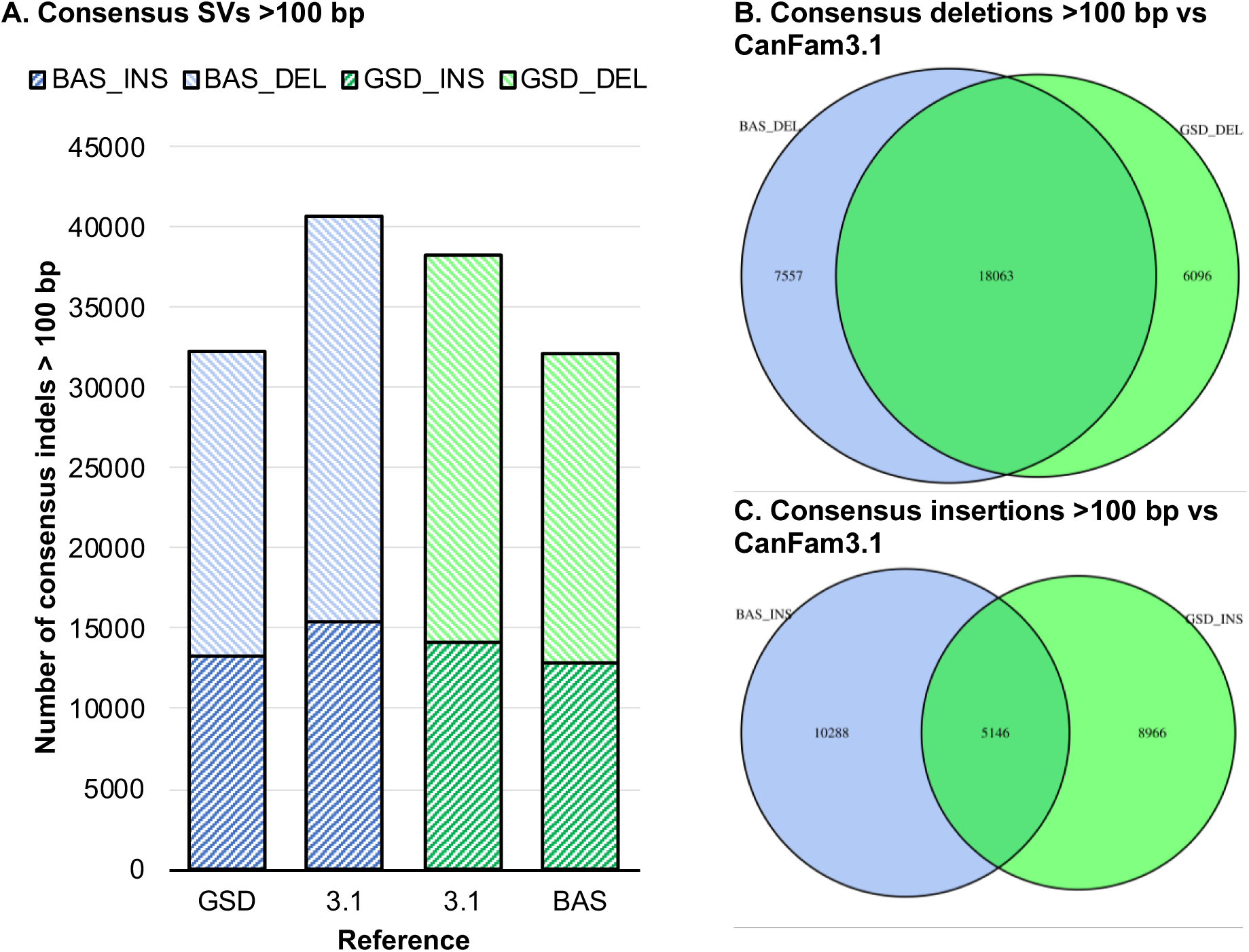
Consensus structural variant calls for combined ONT and PacBio data. High-quality set of consensus structural variant (SV) calls generated from the intersection of the ONT and PacBio SV calls for each breed versus reference comparison, limited to SVs >100 bp long. **A.** Total numbers of SVs called from Basenji CanFam_Bas reads (red) versus CanFam3.1 and CanFam_GSD, and German Shepherd long reads (green) versus CanFam3.1 and CanFam_Bas. **B.** Numbers and overlap of consensus deletion calls for Basenji reads (blue) and GSD reads (green) versus CanFam3.1. **B.** Numbers and overlap of consensus insertion calls for Basenji reads (blue) and GSD reads (green) versus CanFam3.1.

### Short read mapping, SNV / small indel detection

Mapping multiple individuals to a reference genome for variant calling is standard practice in population genomics and is known to be prone to biases in the reference genome. To investigate whether differences identified in variant analyses were due to the Basenji being a basal breed or due to assembly quality difference, short read data from 58 different dog breeds belonging to sixteen different well-supported clades from Parker et al. [10] (Supplementary Table 8, Additional File 2) were mapped on to three reference genomes Basenji (CanFam_BAS), Boxer (CanFam3.1) and GSD (CanFam_GSD). Large-scale structural differences between breeds would be expected to significantly affect read mapping efficiencies for closely-related versus distantly-related breeds, whilst missing assembly data would be expected to result in a systematic reduction in mapping across all breeds. In our analysis, we observe such systematic and breed specific changes in both the number of mapped reads and variants detected.

For the systematic changes, overall trends are exhibited in the total percentage of reads mapped across the three references, with the highest percent of reads mapping to CanFam_GSD, followed closely by CanFam_BAS and the lowest percent of reads mapping to CanFam3.1 (Fig 6A). ANOVA shows these differences are significant (F_2, 171_=819.53, P<0.0001). To investigate this result further and test for interactions we focused upon breeds within each of the monophyletic clades close to or associated with the three reference genomes. Specifically, we included the short-read sequences from the four breeds in the Asian Spitz clade as this close to the basal Basenji lineage. We also included the six breeds European Mastif clade containing the Boxer and three breeds within the New World clade containing the GSD. Overall, the CanFam_Bas and CanFam_GSD performed equally well while the relative mapping was lowest for CanFam3.1 (Fig 6B). Once again ANOVA shows this result is significant (F_8, 30_=32.01, P<0.0001). Next, we considered the capacity of each reference genome to detect SNV’s and indels. In this case CanFam_BAS detected higher number of changes than did either CanFam3.1 or CanFam_GSD (Fig 6C and D; ANOVA F_2, 171_=30.71, and F_2, 171_= 12.08, respectively with P<0.0001 for each). In combination these data attest to the quality of the CanFam_Bas assembly. There is a noticeable depletion of variant calls for the reference breed (Supplementary Fig 6, Additional File 1), but there were no significant interactions between clades and reference genome (data not shown).

**Figure 6.**
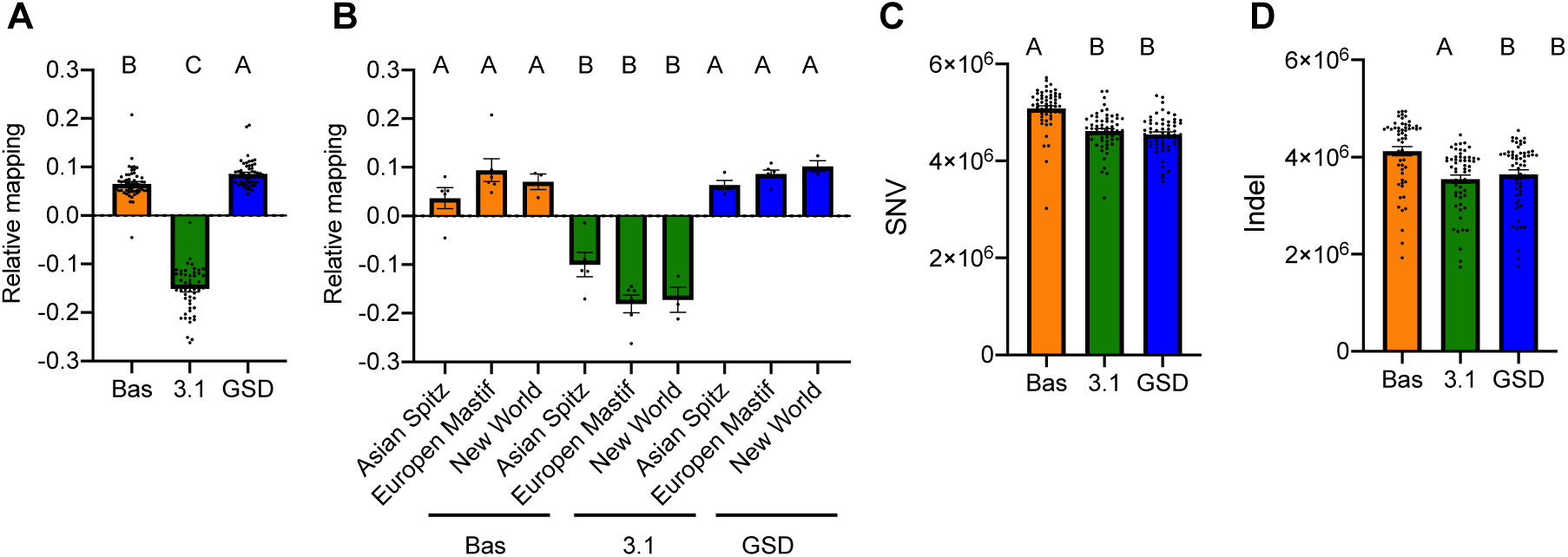
Comparative short read mapping and single nucleotide variant calling for 58 dog breeds versus three reference genomes: CanFam_Bas (Bas), CanFam3.1 (3.1) and CanFam_GSD (GSD). **A.** Relative percentage of reads mapped to each reference calculated by subtracting the mean percentage of reads for the three reference genomes from the number of reads for each reference. **B.** Relative percentage of reads mapped to each reference for three closely related clades determined from Parker et al. [10]. The Basenji is closely related to the Asian Spitz clade. The Boxer is a member of the European Mastiff clade and the GSD is a member of the New world clade. **C.** The number of SNVs. **D.** The number of small indels.

## Discussion

In this manuscript we present a reference-quality assembly of a female Basenji (China), which we designate CanFam_Bas. We also present a second Basenji genome, Wags. The Wags build is of high-quality but not as good as CanFam_Bas, partly because Wags was a male. In total, 64.2% of the missing genes are on the X chromosome, compared to under 16% in the other two individuals. Analysis of long read data mapped onto the CanFam_Bas genome annotation further supports assembly issues as a contributor, with only two autosomal genes lacking any coverage in Wags SMRT data. Equivalently, two other autosomal genes also lacked any coverage in China ONT data, suggesting that even high-quality assemblies can have missing genes in an assembly that may not be biologically lost.

CanFam_Bas is considerably more contiguous and complete than CanFam3.1 (Boxer breed) and slightly more contiguous (Contig N50 37.8 kb vs 20.9 kb) but has slightly lower completeness (BUSCO completeness 92.9% vs 93.0%) than CanFam_GSD [14] (Table 1, Supplementary Table 1, Additional File 2). The ungapped nuclear chromosomes lengths of CanFam_Bas, CanFam3.1, and CanFam_GSD and 2.33 Gb, 2.32 Gb, 2.36 Gb and, respectively. Likely, both assembly quality and breed differences contribute to the 211 missing genes in CanFam3.1. CanFam_GSD had fewer predicted genes missing (95) but a higher percentage (22.0%) also lacking long read coverage. This uncertainty notwithstanding, our analysis highlights the need to consider multiple reference genomes in a pan-genomic approach when a comprehensive analysis is required. These results also highlight the need for considering missing genes carefully on a case-by-case basis as, even with high quality genome assemblies such as those compared in this study, an apparent absence may reflect missing data rather than missing biology.

We noted that the mitochondrial chromosome was missing from the China 1.0 Basenji assembly. An exhaustive search of the genome detected a 33.2kb region consisting of almost two complete copies of the mitochondrial genome on Chromosome 29 that was not present in the other dog assemblies analysed. This assembly error occurred at a real 4,032 bp NUMT. A similar false incorporation of complete mtDNA has been previously reported in the little brown bat [42]. NUMTs are inserted fragments of mtDNA that appear to be present in the nuclear DNA of most, if not all, eukaryotic nuclear genomes [43]. Shotgun genome sequencing data cannot always distinguish NUMTs from mtDNA, which sometimes results in over-stringent removal of NUMTs during genome annotation [43]. Documenting NUMTs in nuclear genome assemblies is important as they have the potential to provide unique insight into population histories and animal well-being [43–45]. Domestic cat nuclear genomes, for example, have 38-76 tandem copies of a 7.9 kb NUMT fragment [46], whilst human NUMTs are polymorphic at 141 loci [43]. NUMT polymorphisms have been used to estimate the age of insertions in human and other primate lineages [44] while five insertions have been implicated in human disease [43, 47]. In total, 212 NUMT blocks (215.1 kb) in 291 fragments (190.5 kb) were detected across the genome, including a previously detected almost full-length (16,577 bp) NUMT on Chromosome 11 [48]. Only one Basenji NUMT was incomplete in the GSD genome. To ease future NUMT comparisons between breeds, we have wrapped up the NUMT discovery and merging methods into an open source tool, NUMTFinder [49].

Sequence analyses estimated six copies of *Amy2B* in China and five in Wags. Previous studies have shown that Basenjis may have 4-18 copies [6], placing these estimates at the lower end of the range. Short read analysis of 11 other Basenji dogs gave similar copy number estimates of four to seven copies, and 4.38 copies in China. This is consistent with the ddPCR estimate of 4.5 copies, although the uneven coverage of short reads is expected to make this less accurate than the long-read approach. Additional work is needed to establish the source of the differences between ddPCR and read depth estimates. The high variation in *Amy2B* copy number suggests at least three possible evolutionary histories of the gene in Basenjis. First, *Amy2B* copies may have differentially accumulated in specific lineages since the divergence from other dog breeds. Second, the ancestral founding population of the modern Basenji may have been polymorphic for *Amy2B.* Third, it remains possible that the *Amy2B* region from other breeds have differentially introgressed into the modern breed dog. Basenji-like dogs are depicted in In Egyptian drawings and models dating back to 1550-1900 BC [11]. Possibly, the Basenji is derived from the Abuwtiyuw, which was a lightly built hunting dog with erect ears and a curly tail that died before 2280BCE [50]. Likely, sequencing the region around *Amy2B* in Basenjis with a higher copy numbers will aid resolving the alternate hypotheses. This large genetic diversity in Basenji *Amy2B* copy has important dietary and veterinary implications. Veterinarians frequently recommend a rice-based diet following complex intestinal surgeries and following diarrhoea because it is thought to be bland and easily digestible. However, dogs with low *Amy2B* copy numbers may not be able to digest rice readily as their serum amylase levels are lower [51].

CanFam_Bas and CanFam_GSD represent two high-quality reference genomes from different breeds. The availability of corresponding Basenji and GSD long read data, provided an excellent opportunity to further investigate the contributions of breed and assembly differences, as real SVs will be represented in the raw read data even if assembly errors have introduced false positives or negatives. Possessing ONT and SMRT data for both GSD and Basenji allows us to overlap the disparate SV call sets to generate a conservative list of SV calls relative to CanFam3.1. This analysis identified over 70,000 SVs in CanFam_Bas relative to CanFam3.1 and over 64,000 SVs in GSD relative to CanFam3.1 (Supplementary Table 7, Additional File 2). There is a high degree of overlap in SVs from GSD and Basenji relative to CanFam3.1 (68% of total basenji SV calls and 77% of GSD SV calls), which highlights potential issues with the current canid reference assembly. Further, each consensus set contains several hundred SVs overlapping annotated exons, highlighting the importance of the selection of appropriate reference genome for analysis of specific genomic regions.

Next, we examined the overlap of the consensus calls for SVs over 100 bp of GSD and Basenji relative to CanFam3.1. We find a high degree of agreement for deletions, with 70.1% of GSD calls and 74.75% of Basenji calls overlapping (Fig 5B), compared to insertions where 33.3% of basenji calls and 36.4% of calls overlap (Fig 5C). Basenji and GSD long reads have 25.9% and 19.1% more SVs called when CanFam3.1 is used as the reference, respectively, and the high degree of overlap is consistent with Boxer-specific SVs and/or assembly issues. GSD-specific and Basenji-specific SV calls are also evident.

To get additional insight into breed differences and the influence of reference genome selection, we mapped short read data from 58 breeds onto CanFam_Bas, CanFam3.1, and CanFam_GSD. If assembly differences are dominated by real differences between breeds then we might expect different breeds to map with different efficiencies onto the three genomes. Differences to due quality, on the other hand, should be reflected across all breeds. With some minor exceptions, short read data from the different breeds consistently mapped better onto CanFam_GSD than CanFam_Bas, which was in turn better than CanFam3.1 (Fig 6A). This is consistent with assembly quality being a dominating factor.

After adjusting for read mapping difference, different reference genomes also produce different SNV and small indel densities for the 58 mapped breeds (Supplementary Fig 6, Additional File 1). Using a Basenji reference genome consistently identifies more variants than either GSD or Boxer (Fig 6, Supplementary Fig 6, Additional File 1). This probably reflects the basal position of Basenji in the breed phylogeny [10]. The basal position of the Basenji makes it useful as a reference for variant analysis as there are clear biases affecting related breeds seen for both the GSD and Boxer reference genomes. On the other hand, CanFam_GSD has slightly higher read mapping levels across breeds, which may provide better total coverage.

Sequencing and assembly efforts are increasingly moving from species reference genomes to breed-specific assemblies, such as those recently published for Great Dane [52], Labrador Retriever [53], and a second GSD [54]. Together, our data suggest that a single high-quality reference should be sufficient for most general analyses, but the generation of breed-specific genomes is likely to be important for canine nutrition and disease studies. The most severe and common ailment in Basenjis is Fancioni Syndrome, in which the renal tubes fail to reabsorb electrolytes and nutrients [55]. Approximately, 10-30% of Basenjis in North America are affected. In 2011, it was shown that Basenji Fanconi Syndrome is caused by a 370bp deletion on canine chromosome 3 [56]. To date no other breeds have been recoded with this same deletion, although other mutations can cause the disease. Likely complex diseases in dogs belonging to different clades may have different underlying causes. For example, it remains unclear whether the same suite of mutations causes hip dysplasia in the GSD (New World clade) and the Labrador (Retriever/ Spaniel clade).

## Conclusions

Here, we present two high quality *de novo* Basenji genome assemblies: CanFam_Bas (China, female) and Wags (male). CanFam_Bas offers improved genome contiguity relative to CanFam3.1 and can serve as a representative basal breed in future canid studies. We generate core genomic information for the Basenji that has the potential to inform future studies of population history and aid disease management. We generate high-quality variants (SNVs, small indels, and SVs) relative to CanFam_Bas, CanFam3.1, and CanFam_GSD. We demonstrate the impact that the reference genome makes on both read mapping and variant detection, illustrating the importance of either selecting the appropriate reference genome or employing a pan-genome approach in future canid studies.

## Methods

### Sequencing and genome assembly of female Basenji, China

#### Sampling

China or Zanzipow Bowies China Girl is an Australian Supreme Champion Kennel Club show dog. Her Australian National Kennel Council registration number is 2100018945. She is bred primarily from Australian lines, with her most recent common ancestor coming from Africa 18 generations previously. She was born on 14 Jan 2016 and is free from all known currently reported diseases.

#### Sequencing

High molecular weight DNA was extracted from 100 µl of blood using the DNeasy Blood and Tissue kit (Qiagen). For long read (Oxford Nanopore) sequencing, 1 µg of DNA was prepared via the Genomic DNA by Ligation kit (SQK-LSK109) as per the manufacturer’s protocol. Prepared DNA (180 ng) was loaded onto a PromethION (FLO-PRO002) flowcell and sequenced with standard parameters. After 48 hours, a nuclease flush was performed, and an additional 115 ng was loaded onto the run. GPU-enabled guppy (v3.0.3) base-calling was performed after sequencing (PromethION high accuracy flip-flop model; config ‘dna_r9.4.1_450bps_hac.cfg’ config).

For short read sequencing whole blood was shipped to BGI, Hong Kong. High molecular weight DNA was extracted, a paired-end library prepared, and the sample run on the BGISEQ-500 sequencing system to generate high quality PE100 data. A total of 767,111,208 clean reads (115.1 Gb) were produced with a lower base call accuracy (Q20) of 95.92%.

#### Assembly

An overview of the China assembly workflow is given in Supplementary Fig 1A, Additional File 1. The ONT reads were assembled with the Flye (v2.6-release) assembler [17, 18]. The resulting contigs were polished with ONT reads using four rounds of Racon (v1.4.3) [19] followed by Medaka (v0.10.0) [20] to minimise error propagation. BGI-seq reads were aligned to the polished assembly with BWA-mem (v 0.7.17) [57] and Pilon (v1.23) (diploid) [21] was used for further error correction. A second assembly was performed using Canu assembler (v1.8.0) [58] and error-corrected with two rounds of Arrow polishing [59]. The Flye assembly was considered more contiguous and therefore selected as the core assembly.

#### Scaffolding

An *in situ* Hi-C library was prepared [24] from a blood sample from China and sequenced to ∼30x coverage (assuming 2.4 Gb genome size). Chromosome-length scaffolding followed the standard DNA zoo methodology (www.dnazoo.org/methods), processing the Hi-C data with Juicer [60] as input for the 3D-DNA pipeline [61]. The resulting candidate scaffolding was manually finished using Juicebox Assembly Tools [22] to produce the final chromosome-length genome assembly. Hi-C matrices are available for browsing at multiple resolutions using Juicebox.js [23] at: https://www.dnazoo.org/post/basenji-the-african-hunting-dog. After scaffolding, all ONT reads were aligned to the assembly with Minimap2 (v2.16) (-ax map-ont) [28] and used by PBJelly (pbsuite v.15.8.24) [25] to fill gaps. BGI reads were re-mapped with BWA-mem (v 0.7.17) and another round of polishing was performed with Pilon (v1.23) (diploid, SNP and indel correction) [21].

#### Final clean-up

The Pilon-polished genome underwent a final scaffold clean-up to generate a high-quality core assembly following Field and colleagues’ processing of Canfam_GSD [14]. Scaffolds were reoriented and assigned to CanFam3.1 chromosomes [15] using PAFScaff (v0.3.0) [14] (Minimap2 v2.16 mapping) [28]. Diploidocus (v0.9.0) [27] was used to screen contamination, remove low-coverage artefacts and haplotig sequences, and annotate remaining scaffolds with potential issues as described in [14]. ONT reads were mapped onto the assembly using Minimap2 (v2.17) (-ax map-ont --secondary=no) [28] and read depth summaries calculated with BBMap (v38.51) pileup.sh [62]. Any scaffolds with median coverage less than three (e.g., less than 50% of the scaffold covered by at least three reads) were filtered out as low-coverage scaffolds. Single-copy read depth was estimated using the modal read depth of 35X across the 5,5578 single copy complete genes identified by BUSCO (v3.0.2b) [33]. This was used to set low-, mid- and high-depth thresholds at 8x, 25x and 68x for PurgeHaplotigs v20190612 [63] (implementing Perl v5.28.0, BEDTools v2.27.1 [64, 65], R v3.5.3, and SAMTools v1.9 [66]). PurgeHaplotig coverage was adjusted to exclude gap regions and scaffolds filtered as in [14] for haplotigs and assembly artefacts (scaffolds with 80%+ bases in the low/haploid coverage bins and 95%+ of their length mapped onto another scaffold by PurgeHaplotigs) or low coverage artefacts (remaining scaffolds with 80%+ low coverage bases). Remaining scaffolds were further classified based on read depth profiles: scaffolds with <20% diploid coverage and 50%+ high coverage were marked as probable collapsed repeats; scaffolds with dominant diploid coverage and >50% match to another scaffold were marked as a possible repeat sequences [14].

#### Genome assembly correction

Main chromosome scaffold integrity was checked using D-GENIES [30] comparisons of different assembly stages with CanFam_GSD chromosomes [14]. Two pairs of fused chromosomes were identified following incorrect joins made by PBJelly (pbsuite v.15.8.24) [25]. Pre-gap-filled HiC scaffolds were mapped onto the assembly using Minimap2 (v2.17) [28] and parsed with GABLAM (v2.30.5) [29] to identify the gap corresponding to the fusion regions. These were manually separated into four individual chromosomes, gap lengths standardised, and scaffolds re-mapped onto CanFam3.1 using PAFScaff (v0.4.0) [26]. Finally, scaffolds under 1 kb were removed following correction of Chromosome 29 (below).

#### Correction of mitochondrial insertion into Chromosome 29

NUMT analysis identified a 33.2 kb region consisting of almost two complete copies of the mitochondrial genome, not present in other dog genome assemblies. GABLAM (v2.30.5) [29] was used to confirm that this region was also absent from the Canu assembly of China v1.1. ONT reads that mapped onto both flanking regions of the 33.2 kb putative NUMT were extracted and reassembled with Flye (v2.7.1) [17, 18]. The new NUMT was approx. 2.8 kb long. To avoid repeated problems with mitochondrial reads mis-polishing the sequence, a subset of 4.82M ONT reads (72.7 Gb, ∼30X) was extracted and mapped onto the assembled region with Minimap (v2.17) [28]. Reads mapping to at least 5 kb of the assembled region including some immediate flanking sequence were extracted (66 reads, 1.50 Mb) and polished with one round of Racon (v1.4.5) [19] (-m 8 -x -6 -g -8 -w 500) and Medaka (v0.7.1) [20] (model r941_prom_high). The polished NUMT region was mapped on to the Chromosome 29 scaffold with GABLAM (v2.30.5) [29] (blast+ v2.9.0 [67] megablast) and stretches of 100% sequence identity identified each side of the NUMT. The mtDNA sequence between these regions of identity was replaced with the re-assembled NUMT sequence.

#### Mitochondrial genome assembly

To assemble the mitochondrion, ONT reads were mapped onto a construct of three tandem copies of the CanFam3.1 mtDNA with minimap2 (v2.17) [28] (-ax map-ont --secondary=no). Reads with hits were extracted using SAMTools (v1.9) [66] fasta and mapped onto a double-copy CanFam3.1 mtDNA with GABLAM (v2.30.5) [29] (blast+ v2.9.0 [67] megablast). “Pure complete” mtDNA ONT reads were identified as those with 99%+ of their length mapping to mtDNA and 99%+ coverage of the mtDNA. These reads were assembled with Flye (v2.7b-b1526) [17, 18] (genome size 16.7 kb) and polished with Racon (v1.4.5) [19] (-m 8 -x -6 -g -8 -w 500) followed by Medaka (v0.7.1) [20] (model r941_prom_high). The polished mtDNA assembly was mapped onto CanFam3.1 mtDNA with GABLAM (v2.30.5) [29] (blast+ v2.9.0 [67] megablast) and circularised by extracting a region from the centre of the assembly corresponding to a single full-length copy with the same start and end positions. Final correction of SNPs and indels was performed by adding the mtDNA to the nuclear assembly, mapping BGI reads with BWA (v0.7.17) and polishing the mtDNA with Pilon (v1.23) [21]. The polished mtDNA was then added back to the nuclear genome for the final *China* assembly.

#### Genome assembly quality assessment

At each stage of the assembly, summary statistics were calculated with SLiMSuite SeqList (v1.45.0) [68, 69], quality was assessed with Merqury (v20200318) (Meryl v20200313, bedtools v2.27.1 [64, 65], SAMTools v1.9 [66], java v8u45, igv v2.8.0) and completeness assessed with BUSCO (v3.0.2b) [33] (BLAST+ v2.2.31 [67], HMMer v3.2.1 [70], Augustus v3.3.2, EMBOSS v6.6.0, laurasiatherian lineage (n=6253)). To account for fluctuations in BUSCO ratings, presence of complete BUSCO genes across assembly stages was also assessed with BUSCOMP (v0.9.4) [71, 72]. Final assembly scaffold statistics and quality assessment was performed with Diploidocus (v0.10.2) (KAT v2.4.2, perl v5.28.0, BEDtools v2.27.1 [64, 65], SAMTools v1.9 [66], purge_haplotigs v20190612, java v8u231-jre, bbmap v38.51, minimap2 v2.17 [28], BLAST+ v2.9.0 [67]). To get a sense of final quality, comparisons were made with the two other dog genomes published at the time of analysis: CanFam3.1 [15] and CanFam_GSD [14].

#### DNA methylation calling

China’s blood DNA methylome libraries were sequenced on the Illumina HiSeq X platform (150 bp, PE), generating 336 million read pairs and yielding 14x sequencing coverage. Sequenced reads were trimmed using Trimmomatic [73] and mapped to the China v1.0 genome reference using WALT [74] with the following parameters: -m 10 - t 24 -N 10000000 -L 2000. The mappability of the MethylC-seq library was 86%. Duplicate reads were removed using Picard Tools (v2.3.0). Genotype and methylation bias correction were performed using MethylDackel with additional parameters: minOppositeDepth 5 -- maxVariantFrac 0.5 --OT 10,140,10,140 --OB 10,140,10,140. The numbers of methylated and unmethylated calls at each CpG site were determined using MethylDackel (https://github.com/dpryan79/MethylDackel). Bisulphite conversion efficiency was 99.71%, estimated using unmethylated lambda phage spike-in control. Nanopore reads were aligned to the generated reference genome using Minimap2 v2.17 [28], and CpG methylation sites were called with f5c v0.6 [75], which is an accelerated version of nanopolish [76]. Methylation frequency was then collated for each CpG site.

#### UMR and LMR calling

Segmentation of basenji’s blood DNA methylome into CpG-rich unmethylated regions (UMRs) and CpG-poor low-methylated regions (LMRs) was performed using MethylSeekR [36] (segmentUMRsLMRs(m=meth, meth.cutoff=0.5, nCpG.cutoff=5, PMDs = NA, num.cores=num.cores, myGenomeSeq=build, seqLengths=seqlengths(build), nCpG.smoothing = 3, minCover = 5).

#### Comparison of DNA methylation calling by PromethION and MethylC-seq

Average MethylC-seq and PromethION DNA methylation for 1 kb genomic bins, UMRs and LMRs were calculated using the *overlapRatios* R function. Scatterplots were generated using *comparisonplot* R function.

### Sequencing and genome assembly of male Basenji, Wags

Wags DNA was derived from blood of a single male. He is registered as American Kennel Club Champion Kibushi the Oracle, born on December 3, 2008. His registration number is HP345321/01. Sire is AM Ch C-Quests Soul Driver, HM827502/02, and his dam is Avongara Luka, HP345312/01, a native female dog imported from the Haut-Ule district of the DRC Congo, 3°24’04.0”N 27°19’04.6”E, in 2006. SMRT sequences for Wags (Fig. 1B) were generated on the Pacific Biosciences Sequel instrument (V2 chemistry) to approximately 45x genome coverage based on a genome size estimate of 2.5 Gb. An overview of the China assembly workflow is given in Supplementary Fig 1B, Additional File 1. All SMRT sequences were assembled with the HGAP4 algorithm, a Falcon based assembly pipeline available through the SMRT Link interphase (SMRT Link v5.0.1.9585) [77]. The assembly was then error corrected with the original SMRT sequences using the Arrow error-correction module [77]. Additional polishing of the assembly for residual indels was done by aligning 32x coverage of Illumina data and the Pilon algorithm [21]. Chromosomal level scaffolds were generated with the same DNA source using the Proximo^TM^ Hi-C genome scaffolding software (Phase Genomics Inc) and finalized by alignment to the CanFam3.1 reference.

### Locus copy number estimation

Copy numbers for specific assembly regions were calculated using Diploidocus (v0.10.0) (runmode=regcnv) [27]. For each animal, long reads were mapped onto the assembly with Minimap2 (v2.17) (no secondary mapping) [28] and the modal read depth across single-copy Complete genes identified by BUSCO v3 [33] (laurasiatherian_db) calculated using SAMTools (v1.9) [66] mpileup. This set the expected single-copy read depth, *X_SC_*. Copy number for a region, *N_reg_* was estimated by dividing the mean read depth across that region, *X_reg_*, by *X_SC_*. The variance and standard deviation of the estimate was calculated using *X_reg_* for all single copy BUSCO genes. For Wags (a male), genes on the X chromosome were excluded from this analysis.

#### Amylase copy number

The copy number of the beta amylase gene *Amy2B* was calculated using Diploidocus (v0.10.0) (runmode=regcnv) [27] using a modification of the single locus copy number estimation (above) to account for multiple copies of the gene in the assembly. First, the AMY2B protein sequence from CanFam3.1 (UniprotKB: J9PAL7) was used as a query and searched against the genome with Exonerate (v2.2.0) [78] to identify assembled copies of the *Amy2B* gene. A full-length (14.8 kb) *Amy2B* gene repeat was also extracted from the CanFam_GSD assembly and mapped onto the assembly with Minimap2 (v2.17) (-x asm5) [28]. Estimated *N_reg_* values were converted into a number of copies by multiplying by the proportion of the query found covered by that region. The total genome *Amy2B* copy number was then calculated as the summed copy number over each hit. To further investigate the robustness of the method and improve the *Amy2B* copy number estimate in CanFam_Bas, analysis was repeated with ONT reads at least 5 kb in length and at least 10 kb in length. These reads should be less susceptible to poor mapping at repeat sequences, but at a cost of reduced coverage. A simpler approach was also applied to short read data for China and eleven Basenji dogs with data available on SRA (Supplementary Table 4, Additional File 2). Reads were mapped onto the China genome with BWA mem [57] and sequencing depth calculated across the annotated AMY2B genes from GeMoMa (Chromosome 6: 46,943,564-46,950,679 and 46,928,705-46,935,822) using samtools v1.11 [66]. The mean sequencing depth per base per AMY2B copy was then compared to the mean sequencing depth across the genome, calculated with BBMap (v38.51) pileup.sh [62]. AMY2B copy number was estimated using the AMY2B:genome read depth ratio.

We also used droplet digital PCR (ddPCR) to directly quantify the *Amy2B* copy number of China DNA [79]. ddPCR was performed using a QX100 ddPCR system (Bio-rad). Each reaction was performed in a 20µl reaction volume containing 10 µl of 2x ddPCR Supermix (Bio-rad), 1µl of each 20x primer/probe, 1µl of DraI restriction enzyme (New England BioLabs #R0129S), 5µl of DNA template (4ng/µl) and 2µl ddH_2_O. Primer sequence for Amy2B: forward 5′-CCAAACCTGGACGGACATCT-3′ and reverse 5′-TATCGTTCGCATTCAAGAGCAA-3′ with FAM probe: 6FAM–TTTGAGTGGCGCTGGG-MGBNFQ. Primer sequence for C7orf28b-3: 5′-GGGAAACTCCACAAGCAATCA-3′ and reverse 5′-GAGCCCATGGAGGAAATCATC-3′ with HEX probe HEX-CACCTGCTAAACAGC-MGBNFQ.

### Nuclear mitochondrial DNA (NUMT) fragment analysis

To make sure that any contiguous NUMTs were identified as a single region, a double-copy CanFam3.1 mtDNA sequence was constructed and then searched against the CanFam_Bas nuclear genome and compressed to unique hits using GABLAM (v2.30.5) [29] (blast+ v2.9.0 [32] blastn, localunique) with a blastn e-value cut-off of 1e-4 [40]. For comparison with published dog NUMTs [48], NUMT fragments with 8 kb were merged into NUMT blocks using NUMTFinder v0.1.0 [49]. Predicted copy number was calculated for each NUMT fragment in China v1.1 using the method described above. Diploidocus (v0.10.0) was also used to calculate number of reads spanning each entire NUMT fragment plus flanking regions of 0 bp, 100 bp, 1 kb and 5 kb. In addition, assembly coverage for each NUMT fragment was calculated for Wags, CanFam3.1 and CanFam_GSD. Each genome was split into 1 Mb tiled fragments and mapped onto CanFam_Bas with Minimap2 (v2.17) [28]. Each BAM file was used for Diploidocus (v0.10.0) regcnv [27] analysis with a single-copy read depth of 1x. Mitochondrial genome coverage was analysed by extracting all 291 NUMT fragment regions with SeqSuite (v1.23.3) [69] and mapping them onto the CanFam3.1 mtDNA chromosome using GABLAM (v2.30.5) [29] (blast+ v2.10 [32] tblastn).

### Genome annotation

Each genome was annotated using GeMoMa [38] (v1.6.2beta, mmseqs2 [80] v5877873cbcd50a6d954607fc2df1210f8c2c3a4b) homology-based gene prediction and nine reference organisms as in Field et al. [14]. To make a fair comparison of the influence of genome quality and completeness on annotation, CanFam3.1 was annotated with the same pipeline. Annotation of CanFam_GSD using the same pipeline was obtained from Field et al. [14].

#### Annotation summary and quality assessment

Annotation summary statistics and the longest protein isoform per gene were generated with SAAGA (v0.4.0) [81]. Annotation completeness was estimated using BUSCO v3 [33] (laurasiatherian, *n*=6253, proteins mode), run on a reduced annotation consisting of the longest protein per gene. To check for truncated or fragmented protein predictions, predicted proteins were mapped onto the Quest For Orthologues reference dog proteome [39] with mmseqs2 v [80]. The best protein hit for each gene was used to calculate a protein length ratio (length of predicted protein / length of reference protein). Percentage coverage of query and hit proteins was calculated with mmseqs2 v [80]. A reciprocal search identified the closest predicted protein for each reference protein. Any reciprocal best hits were marked as predicted orthologues.

#### Annotation copy number and coverage analysis

Predicted copy number was calculated for every protein-coding gene in CanFaBas using the method described above. In addition, assembly coverage for each CanFam_Bas gene was calculated for Wags, CanFam3.1 and CanFam_GSD. Each genome was split into 1 Mb tiled fragments and mapped onto CanFam_Bas with Minimap2 (v2.17) (-ax asm5 -L) [28]. Each BAM file was used for Diploidocus (v0.10.0) regcnv analysis with a single-copy read depth of 1x. In addition, SMRT reads from Wags and ONT reads from the GSD were mapped onto CanFam_Bas with Minimap2 (v2.17) (--secondary=no -L -ax map-pb or -ax map-ont) [28] and the standard predicted copy number calculation applied. Genes with zero coverage were marked 0n. Other genes were binned according to coverage: for mapped assemblies, coverage was rounded to the nearest integer; for long read mapping, coverage was rounded to the nearest 0.5n. Genes with greater than zero but less than 50% coverage were assigned to 0.5n. Any genes exceeding a rounded coverage of 2n were grouped as “3+”.

#### Ribosomal RNA prediction

For each genome, genes for rRNA were predicted with Barrnap (v0.9) [82] (eukaryotic mode, implementing Perl v5.28.0, HMMer v3.2.1 [70] and BEDTools v2.27.1 [64, 65]).

### Whole genome assembly comparisons

Whole genome synteny analysis were performed for the main chromosome scaffolds using the D-GENIES [30] web portal.

### Long read structural variant detection

Structural variant calls were generated using a combination of minimap2 (v2.17-r943-dirty) [28], SAMTools (v1.9) [66], and sniffles (v1.0.11) [83]. In total, four sets of long reads from three samples were analyzed consisting of China the Basenji (Oxford Nanopore), Wags the Basenji (SMRT) and Nala the German Shepherd (Oxford Nanopore and SMRT [14]). Reads were mapped against China v1.0, CanFam3.1 and CanFam_GSD. Analysis was restricted to the main nuclear chromosome scaffolds. Variants for the Basenji and for Nala were annotated with gene model predictions generated using GeMoMa [38] (v1.6.2beta) while CanFam3.1 variants were annotated with Ensembl gene annotations v100 for CanFam3.1 [84]

### Short read mapping, SNV / small indel detection

Representative Illumina data was identified from [85]. All Sequence Read Archive (SRA) IDs associated with the DBVDC bioproject on NCBI (SRP144493) were downloaded along with their metadata and reduced to 126 samples representing the biggest sequencing run (no. bases) per annotated breed. These 126 samples were used for initial read mapping and variant calling (Supplementary Table 9, Additional File 2). Following removal of unknown/mixed/village dog samples, canids other than domestic dogs, and duplicates the remaining breeds were mapped onto those used by Parker et al. [10]. In total, 58 breeds in the significantly monophyletic clades (greater than 70% bootstrap) designated by Parker et al. [10] were considered for analyses (Supplementary Table 8, Additional File 2).

SNVs and small indels were called from the Illumina reads of the 58 representative breeds against three reference genomes (Basenji China v1.0, CanFam3.1, and CanFam_GSD). All Illumina reads were downloaded from the short read archive using the SRA toolkit (v2.10.4) [86]. All samples were analysed using a modified version of an existing variant detection pipeline [87]. Briefly, the pipeline employs BWA (v0.7.17-r1188) for read alignment [88], SAMTools (v1.9) [66] and picard (v2.4.1) (http://broadinstitute.github.io/picard) for binary alignment map (BAM) file preprocessing, and Genome Analysis ToolKit (v3.6) (GATK) for calling SNVs and small indels [89]. The workflow follows GATK best-practices using default parameters throughout. Additionally, SRA reads from each reference genome were aligned using BWA and SAMTools and a consensus variant list generated using an approach described previously [90]. The consensus variant lists for each reference genome were utilized in GATK’s BaseRecalibrator step as the ‘-knownSites’ argument to serve as the required variant truthset. Variant alignment statistics were generated using SAMTools flagstat. Joint variant calls were generated for the larger dataset of 126 samples relative to the three reference genomes and the total number of SNVs and small indels for individual samples tabulated. Variants for the Basenji and CanFam_GSD were annotated with gene model predictions generated using GeMoMa (see above) while CanFam variants were annotated with ENSEMBL gene annotations v100 for CanFam3.1. Read mapping statistics for each sample were calculated using SAMTools to remove secondary mapping and then summarized with BBTools (v38.51) pileup.sh [62]. Numbers were then converted into relative values for each reference by averaging the score for each breed over the three reference genomes and then calculating the difference from the mean. Breeds were considered individually, but mean values for each clade were also calculated. Three clades are of particular importance as they are closely related, or include, the three reference genome assemblies. The Asian Spitz clade is considered closely related to the Basenji. This clade contained the Alaskan Malamute, Shar-Pei, Shiba Inu and Tibetan Mastiff. The European Mastiff Clade contained the Boxer, Bull Terrier, Cane Corseo, Great Dane, Mastiff and Rhodesian Ridgeback. The New World clade contained the Berger Picard, Chinook and German Shepherd. One-way ANOVA’s were employed to detect significant differences between groups. As the same short read samples were examined relative to the three reference genomes statistical significance was set to be P<0.01.

## Abbreviations

BAM: binary alignment map
bp: base pair
ddPCR: digital droplet PCR
CNV: copy number variation
GATK: Genome Analysis Toolkit
Gb: gigabase
GSD: German Shepherd Dog
kb: kilobase
LMR: low-methylated region
Mb: megabase
NGSD: New Guinea Singing Dog
NUMT: nuclear mitochondrial DNA
ONT: Oxford Nanopore Technologies
QFO: Quest For Orthologues
SMRT: Single Molecule Real Time
SNV: single nucleotide variant
SRA: Sequence Read Archive
SV: structural variant
UMR: unmethylated region

## Declarations

### Ethics approval and consent to participate

For China, all experimentation was performed under the approval of the University of New South Wales Ethics Committee (ACEC ID: 18/18B) and with the owner’s written consent. Experiments at the University of Missouri were done with approval from the University of Missouri Animal Care and Use Committee (ACUC 8833) and with owner’s written consent.

## Availability of data and materials

Chromosome-length genome assemblies for both Basenji are available at NCBI (CanFam_Bas (*China*), GCA_013276365.1 (v1.0) and GCA_013276365.2 (v1.2); *Wags*, GCA_004886185.2). The CanFam_Bas mitochondrial genome is available at NCBI Genbank: MW051511. Raw read data is available in the Sequence Read Archive (SRA). SRA identifiers for previously published dog breeds are available in Supplementary Tables 4, 8 and 9 (Additional File 2). Methyl-Seq data is available at GEO accession GSE159396. Supplementary data accompanies this paper at http://www.slimsuite.unsw.edu.au/research/basenji/ and Open Science Foundation (https://osf.io/r3jfm/). Additional datasets used and/or analysed during the current study are available from the corresponding author on reasonable request.

## Authors’ contributions

JWOB coordinated the project. RJE, MAF, and JWOB designed the study. JWOB and WCW funded the project. JWOB and RAZ wrote the ethics approval for the female Basenji. RAZ selected the female China and obtained the sample. JMH performed the DNA extraction and the nuclease flush and directed the ONT data collection for China. JMF and BDR performed the initial China assembly and polishing. AO performed the Hi-C experiment, and OD, RK and ELA conducted the Hi-C analyses. RJE performed the final assembly polishing, curation, correction, analyses of genome completeness, mitochondrial genome assembly, and NUMT analysis. RJE and MAF performed the read-depth gene copy number analyses. SGT completed the digital droplet PCR. KS, JMF and OB conducted the DNA methylation analyses. MAF performed the structural comparisons, dog breed short read mapping and variant calling. MAF, RJE and JWOB performed the dog breed variant analysis. GSJ obtained the ethics approval for the male Basenji, selected the individual Wags and obtained the sample. ER, LH, and WCW performed the Wags assembly. JK performed the genome annotation. RJE, MAF, JMF, BDR, OD, OB and JWOB wrote the manuscript. All authors edited and approved the final manuscript.

## Acknowledgements

We would like to thank Jennifer Power for introducing us to China and Barbara Reisinger, Jon Curby and Jared Reisinger for access to Wags. For China, Brooke Morgan-Burke at the Vineyard Veterinary Hospital provided constant encouragement. ONT data were collected at the Garvan Institute, the BGI-seq data was provided by BGI and the Hi-C data collected at Baylor College of Medicine. For Wags, DNA sequencing was completed at the The Hudson Alpha Institute and assembly at the The McDonnell Genome Institute, Washington University School of Medicine.

## Supplementary Figures and Tables

### Additional file 1 (.pdf): Supplementary Figures

Supplementary Figure 1. The workflow used to construct the de novo genomes.

Supplementary Figure 2. KAT kmer analysis of China assembly.

Supplementary Figure 3. Identification of putative regulatory elements in basenji’s genome using whole-genome bisulphite sequencing.

Supplementary Figure 4. Predicted copy number of BUSCO Complete genes for three dogs based on long-read read depth.

Supplementary Figure 5. CanFam_Bas nuclear mitochondrial DNA (NUMT) coverage.

Supplementary Figure 6. Comparative short read mapping and single nucleotide variant calling for 58 dog breeds versus three reference genomes: CanFam_Bas, CanFam_GSD (GSD) and CanFam3.1 (BOX).

### Additional file 2 (.xlsx): Supplementary Tables

Supplementary Table 1. Assembly statistics for China assembly stages and other dog genomes used in the study.

Supplementary Table 2. Diploidocus statistics and ratings for China scaffolds during tidying.

Supplementary Table 3. GeMoMa summary statistics and annotation comparisons.

Supplementary Table 4. AMY2B short read mapping and estimated copy number in Basenji dogs.

Supplementary Table 5. CanFam_Bas nuclear mitochondrial sequence (NUMT) fragments.

Supplementary Table 6. CanFam_Bas nuclear mitochondrial sequences (NUMTs) merging fragments within 8 kb.

Supplementary Table 7. Structural variant calls for Basenji and German Shepherd long read data versus three reference genomes.

Supplementary Table 8. Dog breeds used for read mapping and SNV analysis.

Supplementary Table 9. SRA runs for representative dog breeds from DBVDC.

