## Additional File 1 for "Chromosome-length genome assembly and structural variations of the primal Basenji dog (*Canis lupus familiaris*) genome"

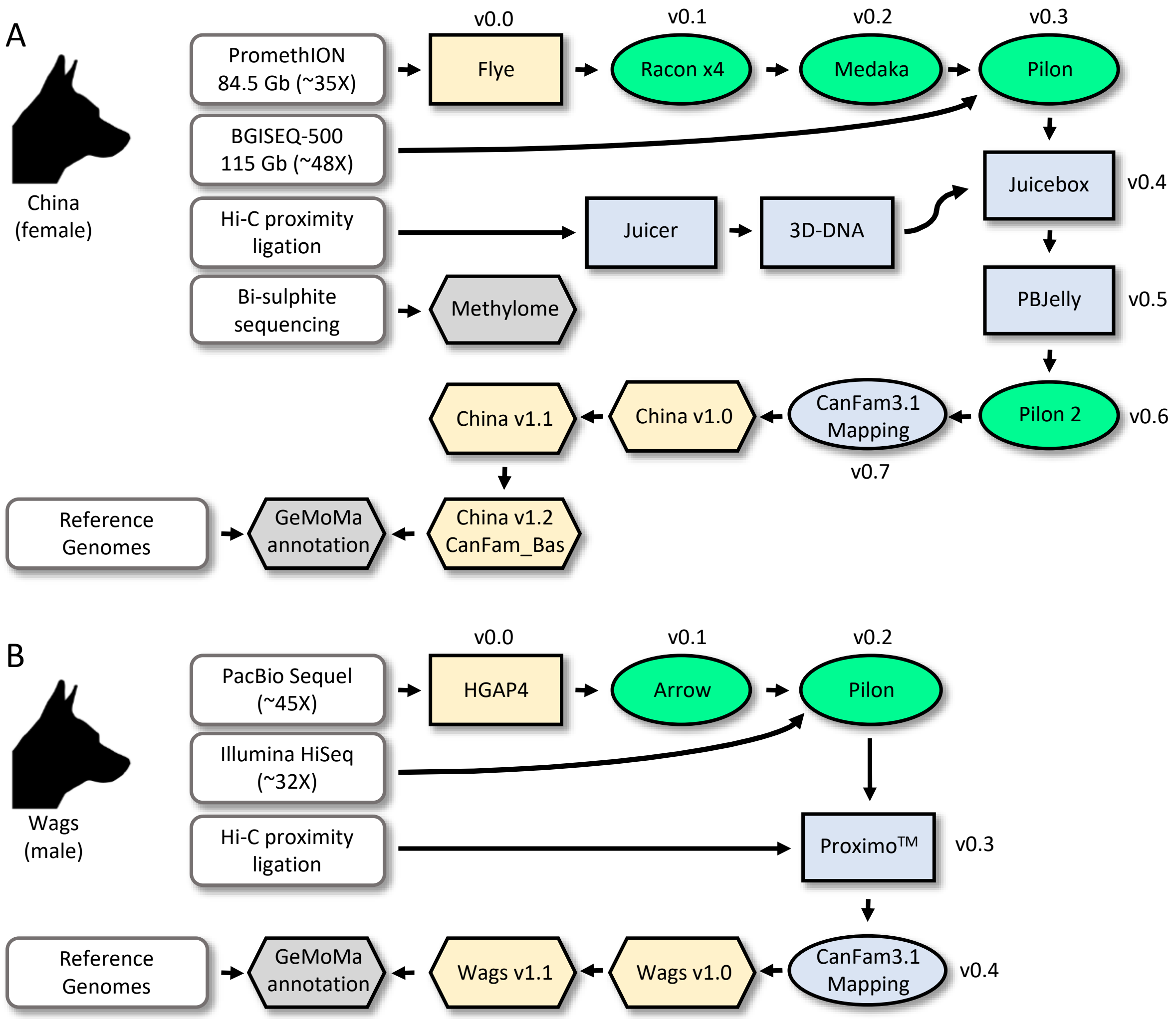


**Supplementary Figure 1. The workflow used to construct the *de novo* genomes.** For both genomes, assembled scaffolds were aligned to CanFam3.1 for chromosome assignments and annotated using the homology-based gene prediction program GeMoMa with 9 reference organisms. See methods for details of software settings and version numbers. Yellow, assembly/curation; Green, polishing; Blue, scaffolding; Grey, other data. **A. China (female).** DNA was derived from from the blood of a single Basenji female, China. Sequences were generated on the Oxford Nanopore PromethION and BGISEQ-500. Long read sequences were assembled with Flye (China v0.0). Long-read error correction was performed four times using Racon (China v0.1) followed by Medaka (China v0.2). Additional short-read error-correction was performed with Pilon (China v0.3). An *in situ* Hi-C library was prepared from the blood of the same individual and processed using Juicer, the 3D-DNA pipeline and Juicebox to produce a chromosome-length genome assembly (China v0.4). The assembly was then long-read gap-filled with the PBJelly algorithm (China v0.5), and polished for a second time using Pilon (China v0.6). Scaffold were mapped onto CamFam3.1 for chromosome assignments (China v0.7) before a final tidying step with Diploidocus to produce China v1.0. Subsequent error-correction of fused chromosomes (China v1.1) and mtDNA insertion (China v1.2) produced the final CanFam_Bas reference genome. **B. Wags (male).** DNA was derived from blood of a single Basenji male, Wags. Sequences were generated on the Pacific Biosciences Sequel instrument. All SMRT sequences were assembled with the HGAP4 algorithm (Wags v0.0) then error corrected using the Arrow error-correction module (Wags v0.1). Additional polishing of the assembly for residual indels was done by aligning 32x coverage of Illumina data and the Pilon algorithm (Wags v0.2). Chromosomal level scaffolds were generated with the same DNA source using the ProximoTM Hi-C genome scaffolding software (Phase Genomics Inc) (Wags v0.3) and finalized by alignment to the CanFam3.1 reference (Wags v0.4). Additional curation produced the final Wags genome (Wags v1.1) used in this study. *Basenji silhouette credit: Arran Morton.*

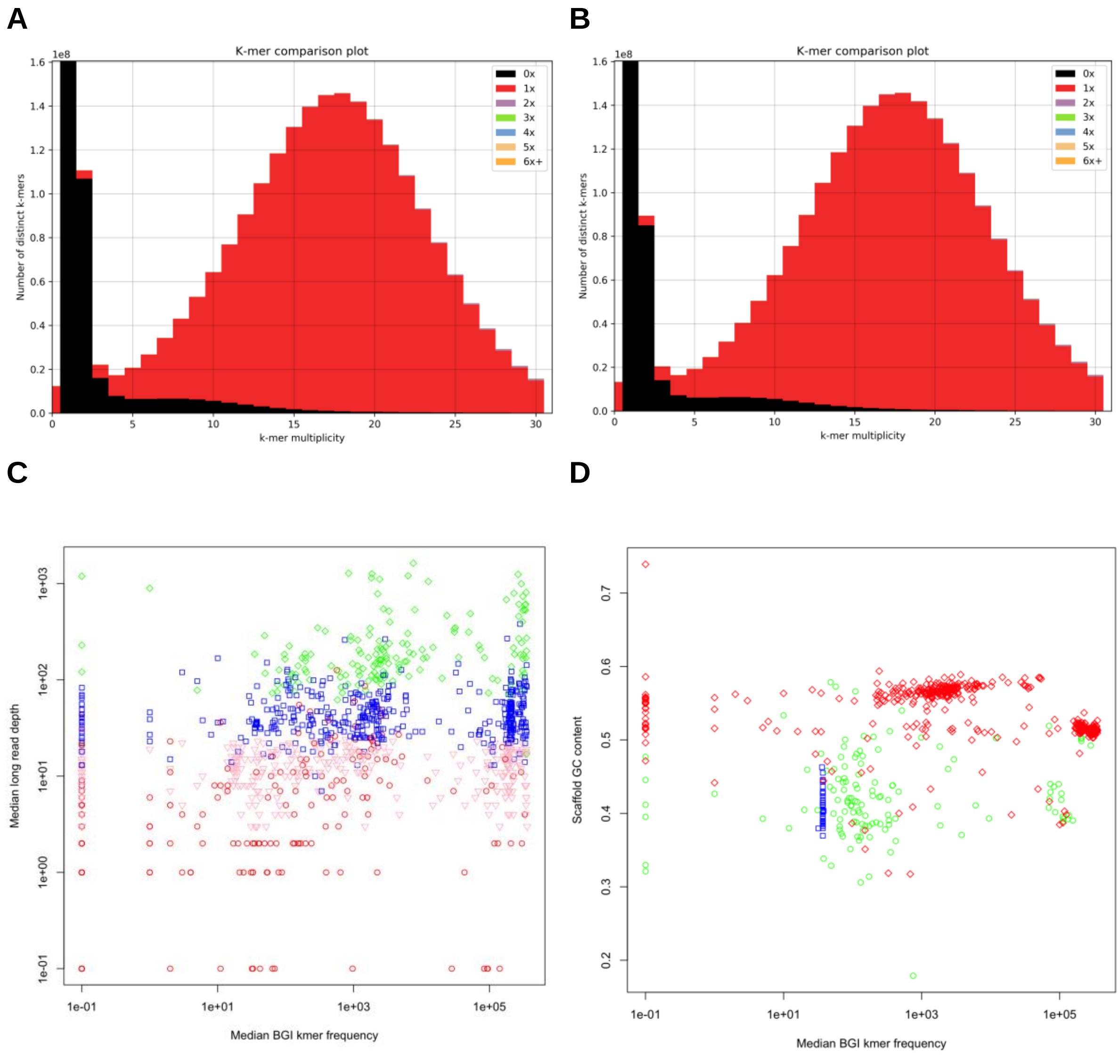


**Supplementary Figure 2. KAT kmer analysis of China assembly.** BGI read kmer frequency distributions for kmers with different assembly copy numbers derived from **(A)** Read 1 and **(B)** Read 2. **C.** Median long read depth versus median 10x kmer frequency and Diploidocus ratings for all scaffolds. Blue square, keep; Green diamond, repeat; Pink triangle, quarantine; Red circle, purge. **D.** Scaffold GC content versus median 10x kmer frequency for tidied assembly. Blue square, chromosome; Green circle, placed (unlocalised) scaffold; Red diamond, unplaced scaffold.

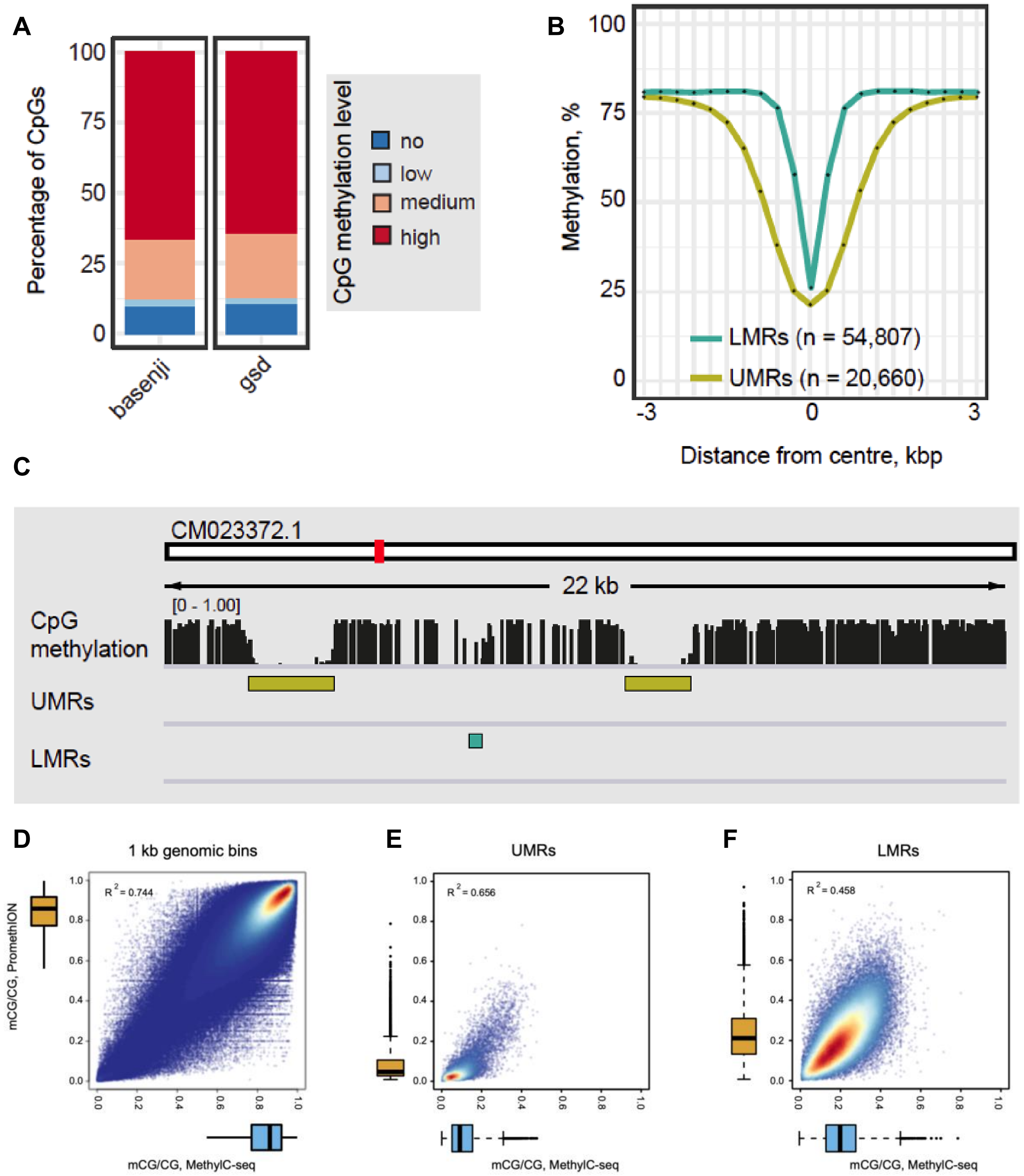


**Supplementary Figure 3. Identification of putative regulatory elements in basenji's genome using whole-genome bisulphite sequencing and comparison of DNA methylation calling by PromethION and MethylC-seq. A.** Percentage of CpG sites with different levels of methylation in Basenji and German Shepherd Dog blood. Levels of methylation are as follows: high (80-100%), medium (20-80%), low (>0-20%), no (0%). **B.** Average CpG methylation profiles at UMRs (n = 20,660) and LMRs (n = 54,807). **C.** IGV browser tracks showing CpG methylation profile of basenji's blood DNA as well as examples of UMRs and LMRs. **D-E.** Scatterplots showing the correlation between the average CpG methylation of 1 kbp genomic bins (**D**), UMRs (**E**) and LMRs (**F**) detected by MethylC-seq (x-axis) and PromethION (y-axis). Each dot represents a single genomic region. The density of data points goes from blue (low) to red (high). Average methylation Adjusted $R^2$ values are indicated on each scatterplot.

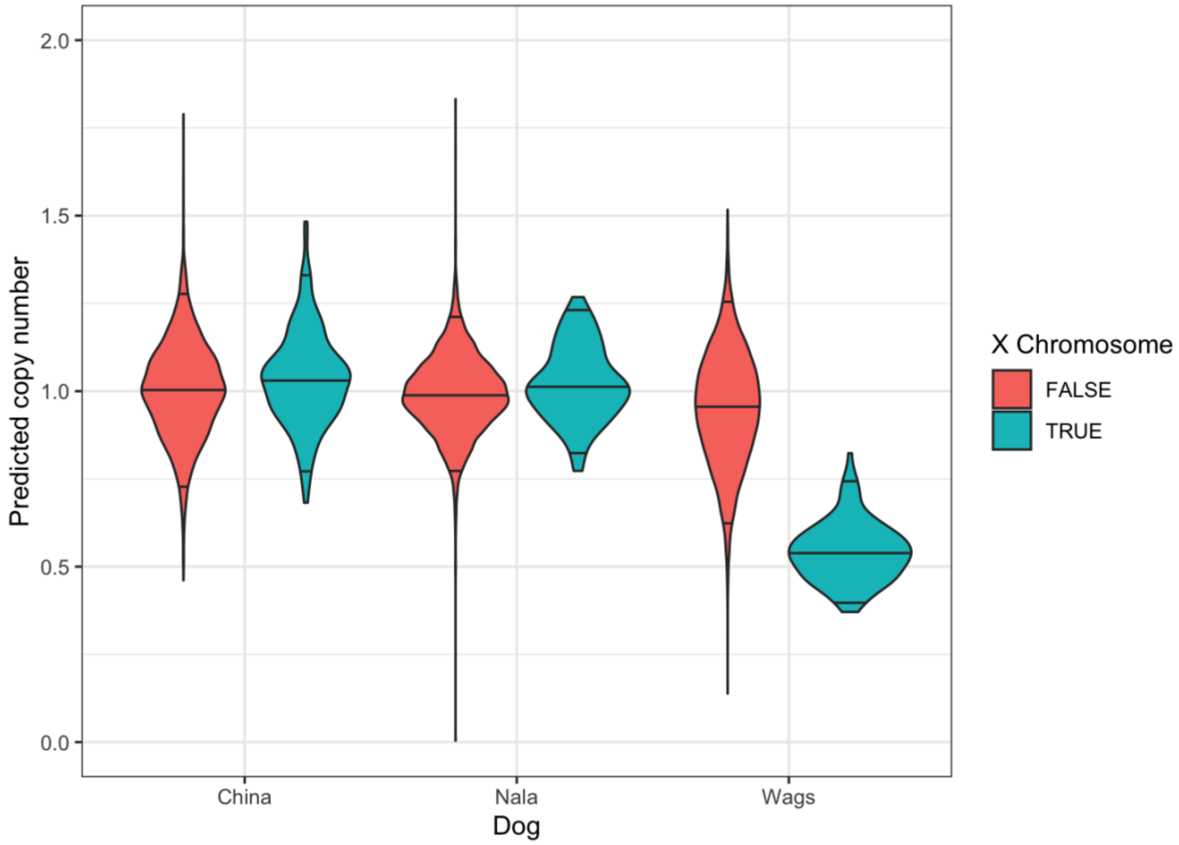


**Supplementary Figure 4. Predicted copy number of BUSCO Complete genes for three dogs based on long-read read depth.** BUSCO Complete genes for CanFam_Bas ("China"), CanFam_GSD ("Nala") and Wags were used to establish the dominant single copy read depth. Mean coverage across each gene was divided by single copy depth to estimate the predicted copy number distribution for autosomes (red) and the X-chromosome (green).

A

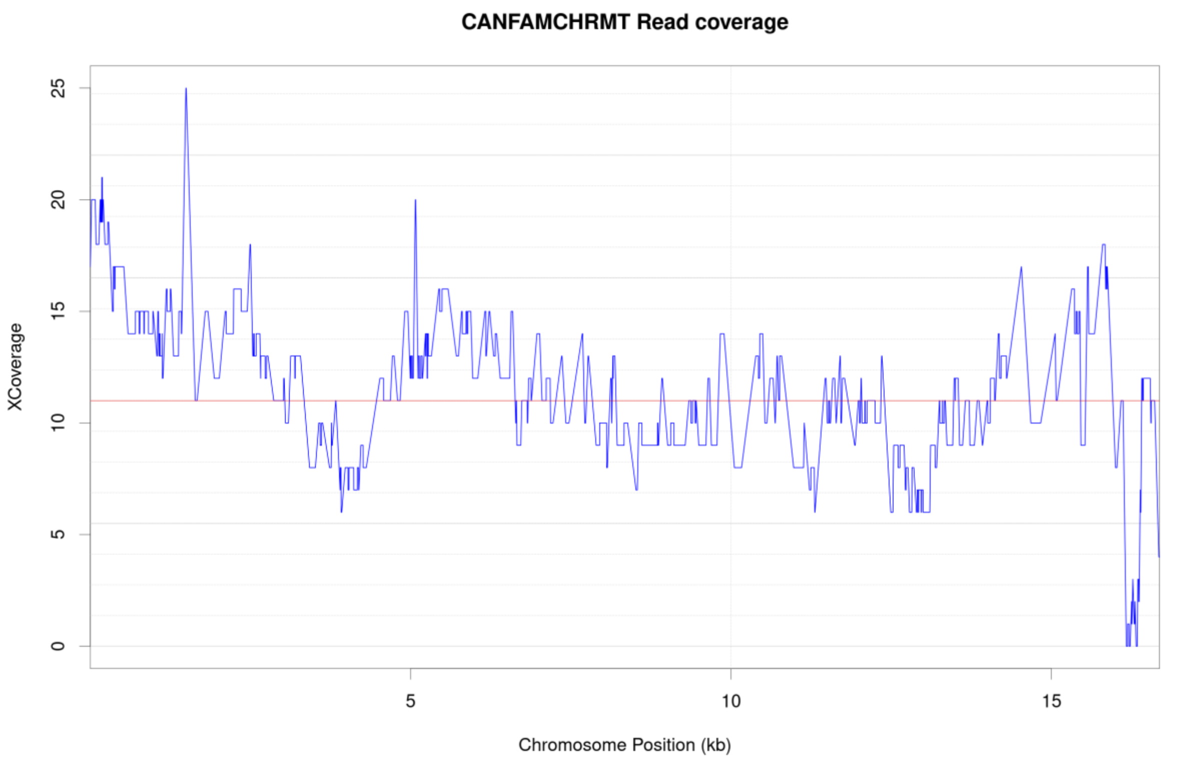


B

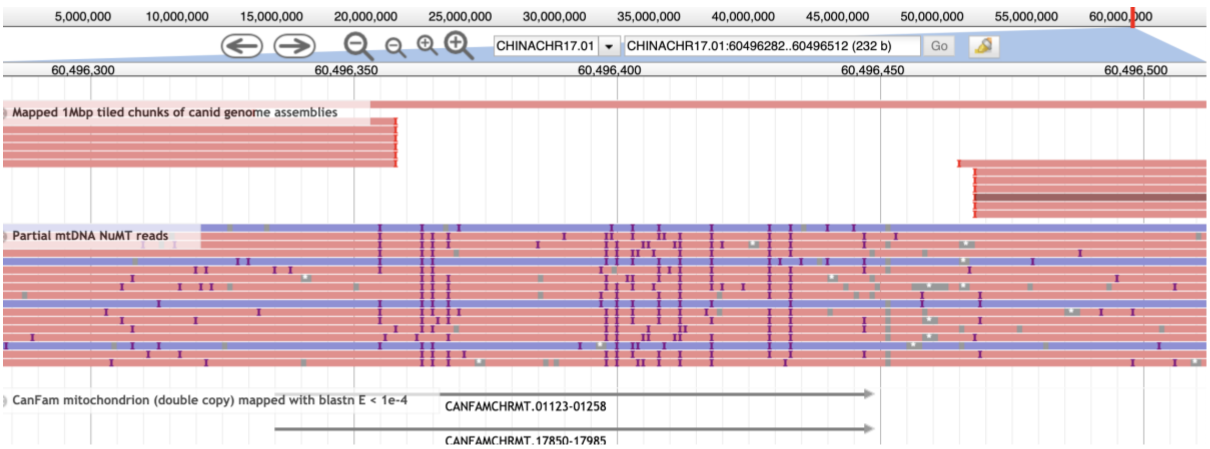


**Supplementary Figure 5. CanFam_Bas nuclear mitochondrial DNA (NUMT) coverage. A.** Depth of coverage for CanFam_Bas NUMT sequences mapped onto the CanFam3.1 mitochondrial genome. Median coverage of 11X is marked as a red line. **B.** Screenshot of WebApollo browser for the NUMT on CanFam_Bas chromosome 17 that was not full-length in CanFam_GSD. The top track shows dog genomes mapped onto CanFam_Bas. The contiguous mapping is CanFam_Bas itself, with all other genomes having a gap in this region, spanning most of the NUMT and approx. 20 bp downstream. The middle track shows CanFam_Bas raw long-read data, clearly spanning the full NUMT, which is shown in the bottom track.

### A. Single nucleotide variants per kilobase of reference with read coverage

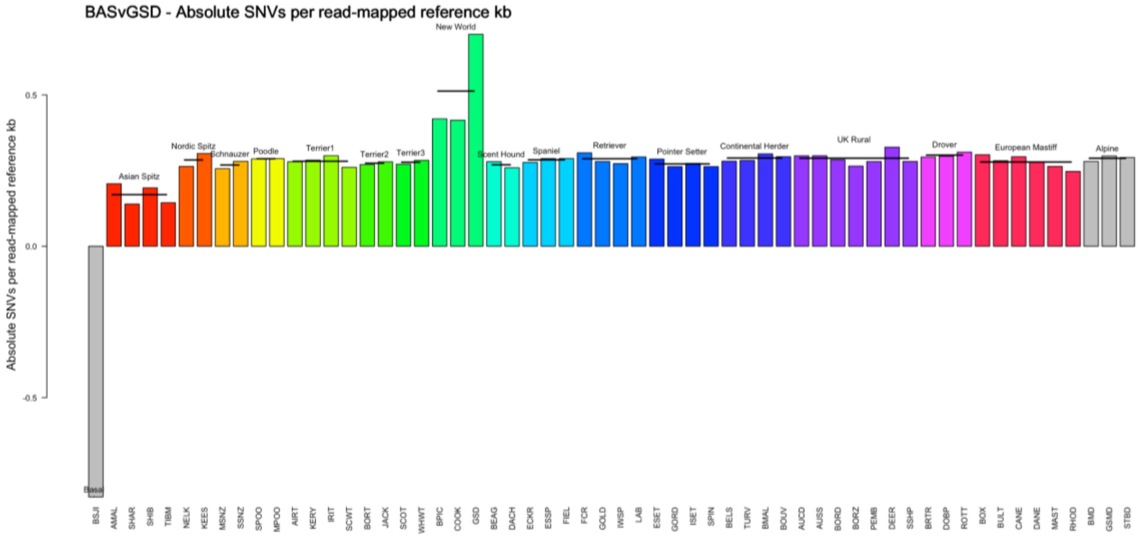


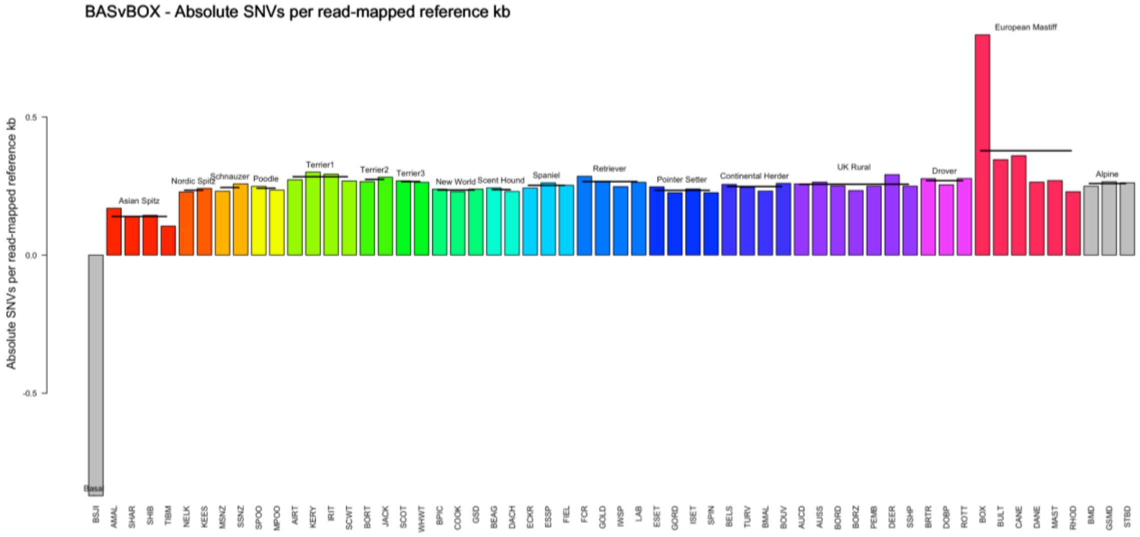


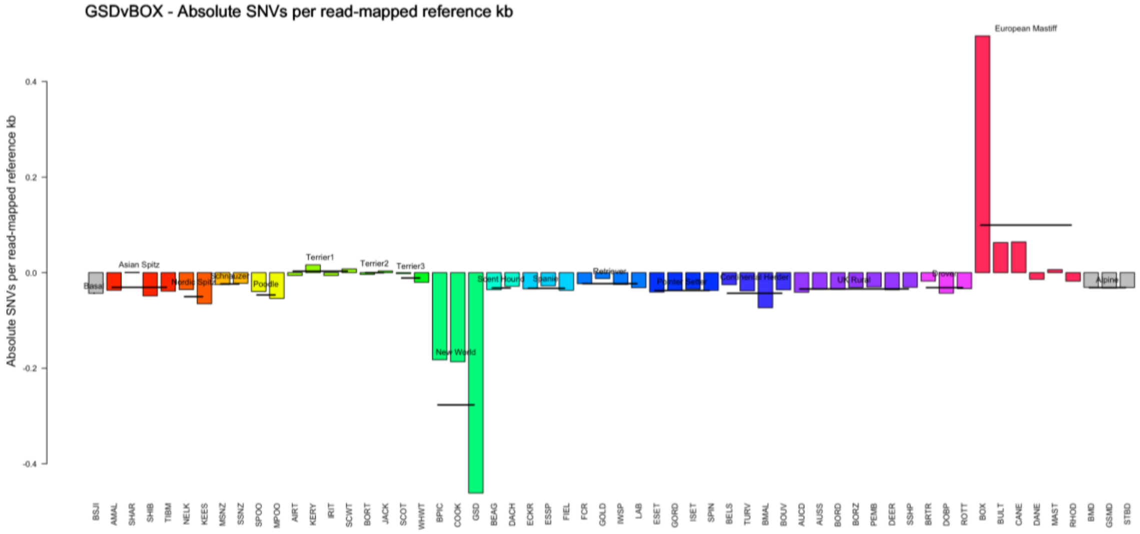


Cntd. on next page

### B. Small indels per kilobase of reference with read coverage

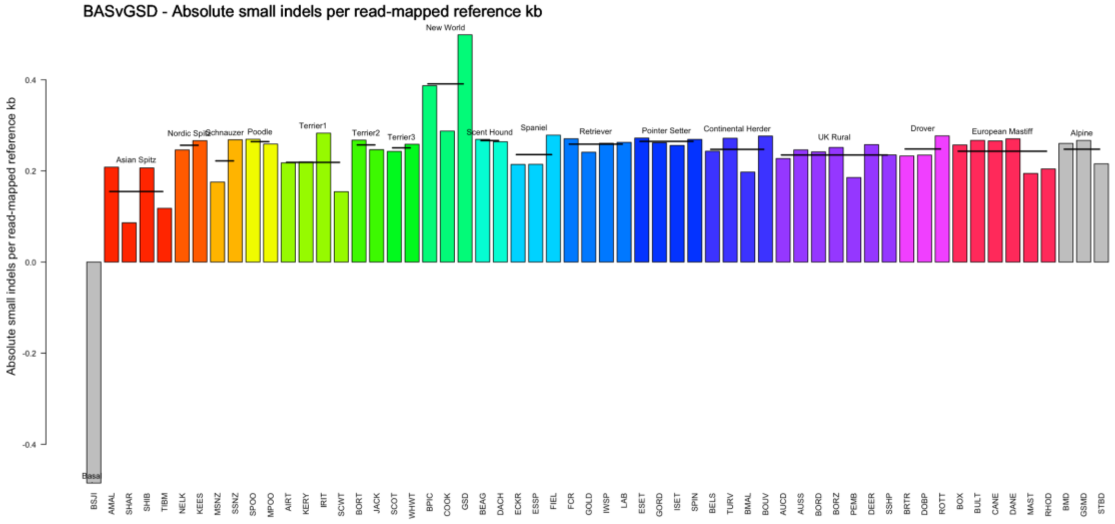


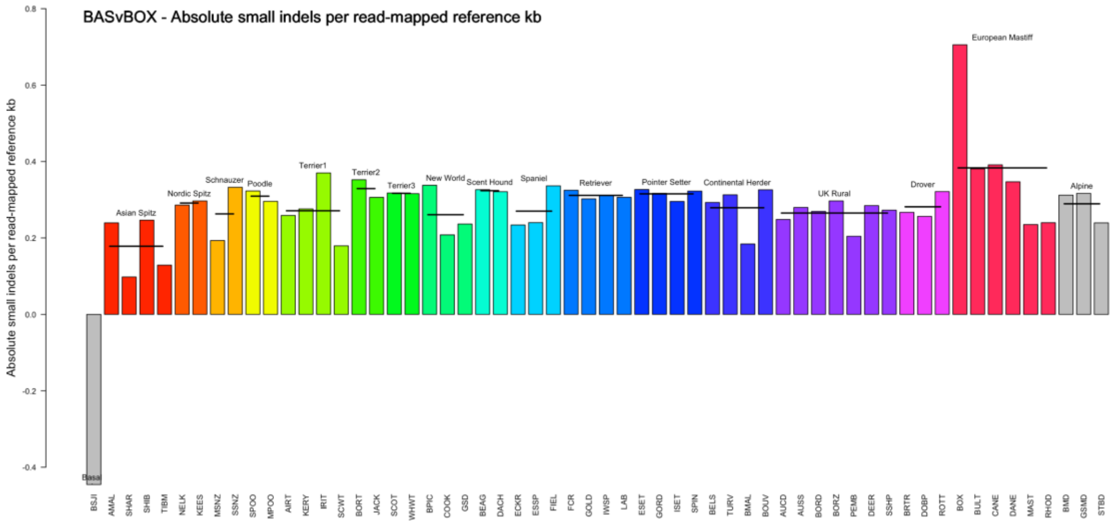


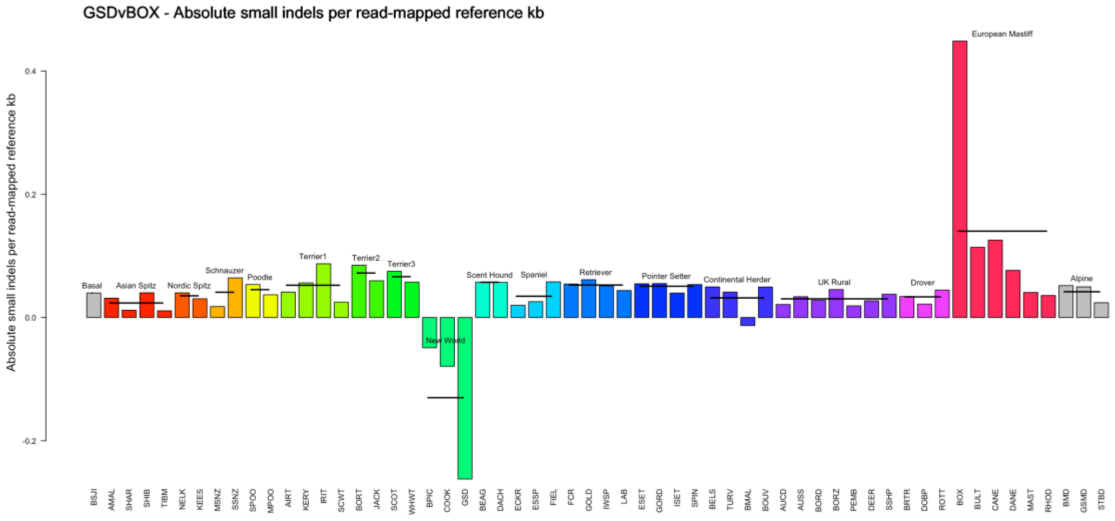


Legend on next page

**Supplementary Figure 6. Comparative short read mapping and single nucleotide variant calling for 58 dog breeds versus three reference genomes: CanFam_Bas, CanFam_GSD (GSD) and CanFam3.1 (BOX).** Each panel shows three comparisons showing the difference between results from a pair of reference genomes: top, CanFam_Bas - CanFam_GSD; middle, CanFam_Bas - CanFam3.1; bottom, CanFam_GSD - CanFam3.1. Each bar represents a sample from a different dog breed, coloured and grouped by well-supported clades from Parker et al. [13] and labelled with a three letter abbreviation for that breed (see Methods and Supplementary Table 6 for details). The mean for each clade is shown as a thick black line. **A.** The number of single nucleotide variants per kilobase, adjusted for mapped read coverage of the reference genome. **B.** The number of small indels per kilobase, adjusted for mapped read coverage of the reference genome.
